# Phosphatidic acid is an endogenous negative regulator of PIEZO2 channels and mechanical sensitivity

**DOI:** 10.1101/2024.03.01.582964

**Authors:** Matthew Gabrielle, Yevgen Yudin, Yujue Wang, Xiaoyang Su, Tibor Rohacs

**Author notes:** School of Pharmaceutical Sciences, Tsinghua-Peking Center for Life Sciences, Beijing Frontier Research Center for Biological Structure, Tsinghua University, Beijing, China (Yujue Wang).

## Abstract

Mechanosensitive PIEZO2 ion channels play roles in touch, proprioception, and inflammatory pain. Currently, there are no small molecule inhibitors that selectively inhibit PIEZO2 over PIEZO1. The TMEM120A protein was shown to inhibit PIEZO2 while leaving PIEZO1 unaffected. Here we find that TMEM120A expression elevates cellular levels of phosphatidic acid and lysophosphatidic acid (LPA), aligning with its structural resemblance to lipid-modifying enzymes. Intracellular application of phosphatidic acid or LPA inhibited PIEZO2, but not PIEZO1 activity. Extended extracellular exposure to the non-hydrolyzable phosphatidic acid and LPA analogue carbocyclic phosphatidic acid (ccPA) also inhibited PIEZO2. Optogenetic activation of phospholipase D (PLD), a signaling enzyme that generates phosphatidic acid, inhibited PIEZO2, but not PIEZO1. Conversely, inhibiting PLD led to increased PIEZO2 activity and increased mechanical sensitivity in mice in behavioral experiments. These findings unveil lipid regulators that selectively target PIEZO2 over PIEZO1, and identify the PLD pathway as a regulator of PIEZO2 activity.

## INTRODUCTION

PIEZO1 and PIEZO2 are non-selective cation channels activated by mechanical force^1^. PIEZO2 is expressed in peripheral sensory neurons of the dorsal root ganglia (DRG), and it is indispensable for gentle touch and proprioception in mice and in humans, and has been implicated in injury-induced mechanical pain^2^. PIEZO1 has a broader expression profile, and is involved in a variety of processes, including blood and lymphatic vessel formation, red blood cell volume regulation, and epithelial cell division^3^.

Both PIEZO1 and PIEZO2 channels are inhibited by non-specific blockers of mechanically-activated ion channels such as gadolinium, ruthenium red and GsMTx4^4^. Screening small molecule libraries resulted in the discovery of Yoda1, as well as Jedi1 and Jedi2, selective activators of PIEZO1 channels^5,6^, but the same screens did not identify chemical activators of PIEZO2. Chemical modification of Yoda1, resulted in Dooku^7^, a compound that antagonizes Yoda1 activation of PIEZO1. Currently there are no known endogenous nor exogenous inhibitors that selectively inhibit PIEZO2 over PIEZO1. Selective PIEZO2 inhibitors would be highly valuable research tools, and would have substantial therapeutic potential against mechanical pain^2,4^.

We have recently shown that coexpression of the TMEM120A (a.k.a. TACAN) protein robustly inhibited PIEZO2, but not PIEZO1 channel activity^8^. TMEM120A was proposed earlier to be an ion channel involved in sensing mechanical pain^9^, but subsequent studies raised doubts about its ability to function as an ion channel and respond to mechanical stimuli^8,10–13^. Structural studies revealed that TMEM120A is a homodimer of two subunits with six transmembrane domains in each^10,12,13^ with structural homology to the long chain fatty acid elongase enzyme ELOVL7^14^. Consistent with its potential function as a lipid modifying enzyme, the TMEM120A structures contained a coenzyme A (CoA) molecule^10,12,13^. The specific enzymatic activity of TMEM120A, however, is unknown^10^, see our recent review for details^15^.

Here, we hypothesized that TMEM120A inhibited PIEZO2 currents by altering cellular lipid content and aimed to identify the lipid regulator that selectively inhibits PIEZO2 without affecting PIEZO1 activity. To achieve this, we conducted liquid chromatography tandem mass spectrometry (LC-MS/MS) experiments, which revealed that TMEM120A expression robustly elevates phosphatidic acid and lysophosphatidic acid (LPA) levels with saturated acyl chains. Furthermore, we observed that mutating a putative catalytic residue in TMEM120A reduced its inhibitory effect. Intracellular delivery of phosphatidic acid, or LPA through the whole-cell patch pipette inhibited PIEZO2 activity, while leaving PIEZO1 unaffected. Long-term incubation with ccPA, a metabolically inert analogue of phosphatidic acid and LPA, effectively inhibited heterologously expressed PIEZO2 channels, and decreased both the proportion and amplitudes of rapidly adapting mechanically-activated currents in DRG neurons. Optogenetic activation of phospholipase D (PLD), a signaling enzyme that generates phosphatidic acid, inhibited PIEZO2, but not PIEZO1 channel activity. Inhibition of PLD enzymes potentiated both heterologously expressed and native PIEZO2, while not affecting PIEZO1 activity. PLD inhibition also increased mechanical sensitivity in mice in behavioral experiments. We identify phosphatidic acid and LPA as specific inhibitors of PIEZO2 channels, and our data indicate that PLD enzymes, which produce phosphatidic acid, modulate PIEZO2 activity. These findings unveil lipid regulators that selectively target PIEZO2 over PIEZO1, and hold promise for the development of specific PIEZO2 inhibitors.

## RESULTS

### Changes in cellular lipid content mediate inhibition of PIEZO2 by TMEM120A

Earlier research indicates that expression of TMEM120A results in robust inhibition of PIEZO2, but not PIEZO1 channels^8^. TMEM120A shows structural homology to the long chain fatty acid elongase enzyme ELOVL7^10,12,13^, but its potential lipid-modifying enzymatic activity has not yet been determined^15^. To test the hypothesis that TMEM120A inhibits PIEZO2 activity by modifying cellular lipid content, we performed LC-MS/MS experiments in cells transfected with TMEM120A, and compared them to mock (vector) transfected cells, and cells transfected with TMEM120B, a homologue of TMEM120A that did not inhibit PIEZO2 activity^8^. We used Neuro2A (N2A) cells in which endogenous PIEZO1 was deleted using CRISPR (N2A-Pz1-KO)^16,17^. We found this cell line to be a good model system to study heterologously expressed PIEZO2 channels^8^, and also used it for all electrophysiology experiments in this study. Cells transfected with TMEM120A showed a large, 4-fold increase in the levels of phosphatidic acid and lysophosphatidic acid (LPA) with fully saturated acyl chains (**Fig. 1a**). Diacylglycerol (DAG) showed a small, ∼20% increase, and triacylglycerol (TG), showed a 1.9-fold increase compared to control, mock transfected cells (**Fig. 1a**). Considering that transfection efficiency was ∼40%, these bulk changes likely underestimate the changes at the individual cell level. Cells transfected with TMEM120B did not show these changes, except for a small, ∼30% increase in TG levels (**Fig. 1a**). These lipids are intermediates in the Kennedy pathway of TG synthesis from glycerol-3 phosphate (**Fig. 1b**), and they were largely restricted to lipid species with fully saturated acyl chains. Lipids with one or more double bonds in their acyl chains showed either no, or smaller changes (**Supplementary Fig. 1**).

**Figure 1.**
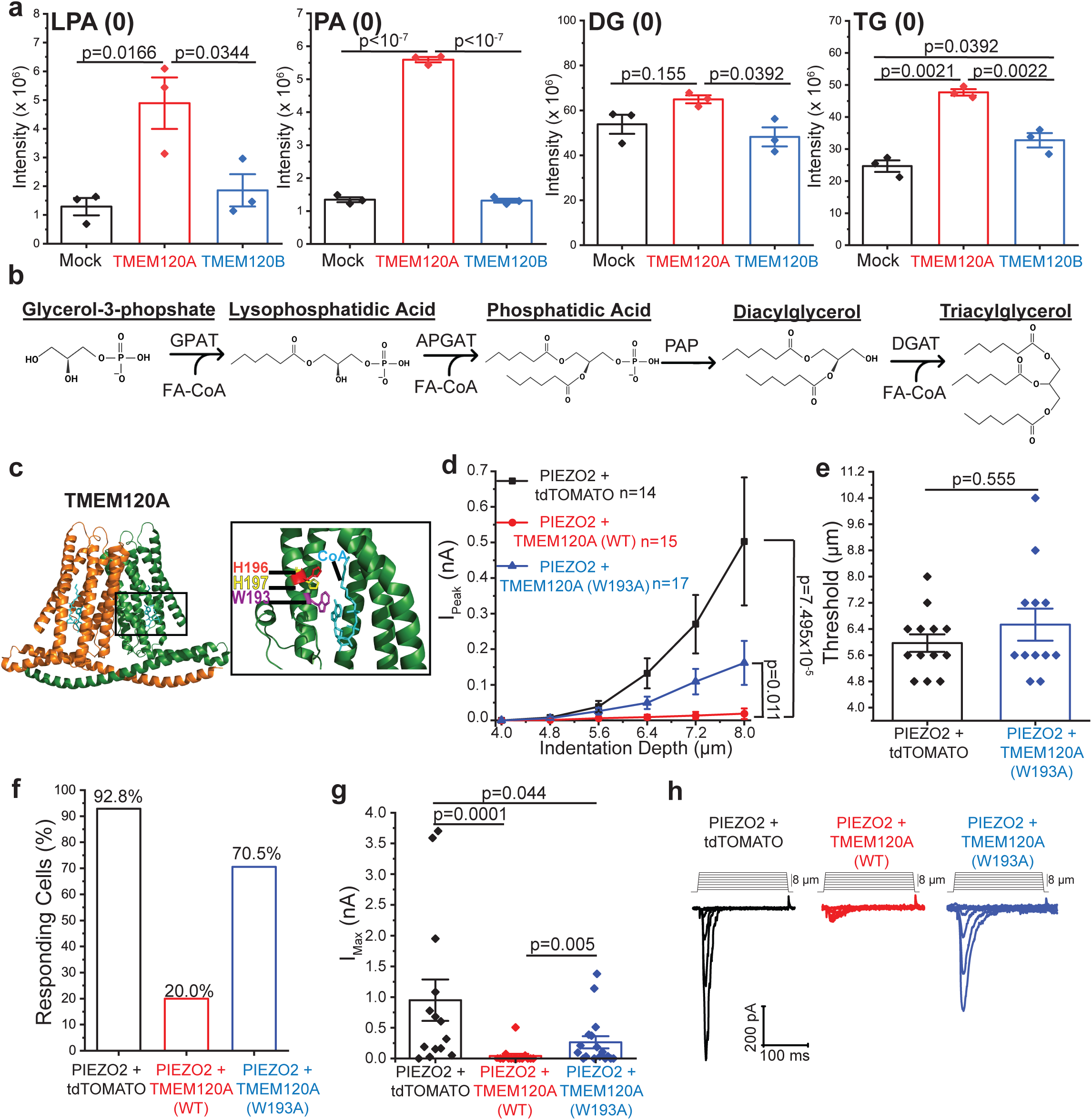
Changes in cellular lipid content mediate inhibition of PIEZO2 by TMEM120A. **(a)** LC-MS/MS experiments using N2A-Pz1-KO cells transiently transfected with vector (mock, black), TMEM120A (red), or TMEM120B (blue) as described in the methods section. Scatter plots and mean ± SEM of the relative saturated lipid intensities detected for n=3 independent transfections for LPA (ANOVA, F(2,6)=9.274, p=0.0146), phosphatidic acid (PA) (ANOVA, F(2,6)=1148, p 1.77x10^-8^), DG (ANOVA, F(2,6)=5.528, p=0.0435), TG (ANOVA, F(2,6)=44.500, p=0.0002). P values for post-hoc Tukey tests are displayed on plots. **(b)** Scheme of the Kennedy pathway for de novo TG synthesis. Lipids are depicted with short acyl chains because of spatial restrictions. **(c)** Structure of TMEM120A with magnified (black box) of residues interacting with CoA generated using PyMOL from the publicly available pdb file 7F3T^12^. **(d)** Whole-cell voltage clamp experiments at -60 mV in N2A-Pz1-KO cells transiently transfected with PIEZO2-GFP, and tdTOMATO (black), TMEM120A-tdTom-WT (red), or TMEM120A-tdTom-W193A (blue). Mechanically-activated currents were evoked by indentations of increasing depth with a blunt glass probe. Current amplitudes are plotted (mean ± SEM) and statistical difference for the area under curve (AUC) for 4.0-8.0 µm stimuli calculated with the Mann-Whitney test. **(e)** Scatter and mean ± SEM for the threshold of membrane indentation to elicit mechanically-activated current (Mann-Whitney). **(f)** Percent of cells displaying mechanically-activated currents. **(g)** Scatter and mean ± SEM for maximum currents (Kruskal-Wallis, χ^2^=17.467, df=2, p=0.0001; Mann-Whitney tests displayed). **(h)** Representative current traces.

Next, we mutated a putative catalytic residue that interacts with CoA in the TMEM120A structures (**Fig. 1c**), and we found that co-expressing this TMEM120A-W193A mutant induced a significantly smaller inhibition of PIEZO2 than the wild-type TMEM120A (**Fig. 1d-h**). We obtained similar results with a triple mutant of TMEM120A, in which we introduced two additional mutations in two adjacent histidine residues that were implicated in catalytic activity of ELOVL^10^ (W193A-H196A-H197A; AxxAA) (**Supplementary Fig. 2a-d**). In these experiments, we used TMEM120A, and its mutants tagged with the red fluorescent protein tdTomato, and PIEZO2 tagged with EGFP. **Supplementary Fig. 2e-g** shows that tdTomato fluorescence in total internal reflection fluorescence (TIRF) experiments was not different in cells transfected with wild-type or W193A or AxxAA mutants of TMEM120A. TIRF microscopy shows fluorescence in the plasma membrane and a narrow sub-plasmalemmal region, thus these experiments indicate that the reduced inhibition by the mutant TMEM120A was not caused by reduced surface expression. EGFP fluorescence in the TIRF mode was also not different between cells transfected with PIEZO2-GFP and the mutant and wild type TMEM120A or tdTomato, indicating that inhibition of PIEZO2 by TMEM120A was not caused by reduced surface expression, a finding consistent with our earlier result^8^.

We also found that inhibiting GPAT, the first enzyme in the Kennedy pathway (**Supplementary Fig. 2h**) significantly reduced the inhibition of PIEZO2 currents by TMEM120A (**Supplementary Fig. 2i-m**). The reduction of inhibition was small, which is likely due to an alternative pathway to generate LPA from glycerol-3-phosphate by the DHAPAT and Acyl-DHAP Reductase enzymes (**Supplementary Fig. 2h**). Overall, these data indicate that the changes in lipid composition induced by TMEM120A play a role in the inhibition of PIEZO2.

### Phosphatidic acid and LPA inhibit PIEZO2 but not PIEZO1 channels

To identify the lipid molecule(s) that inhibit PIEZO2 activity, we performed whole-cell patch clamp experiments in N2A-Pz1-KO cells transfected with PIEZO2, or PIEZO1, and evoked mechanically-activated inward currents using increasing indentations with a blunt glass probe (**Fig. 2**). Consistent with earlier results^16,18,19^ non-transfected N2A-Pz1-KO cells did not display mechanically-activated currents in response to indentations with the blunt glass probe (data not shown), but cells expressing PIEZO2 (**Fig. 2a,d**) or PIEZO1 (**Fig. 2e,h**) showed characteristic rapidly inactivating mechanically-activated currents. We delivered various lipids to the intracellular compartment through the whole cell patch pipette, and compared mechanically-activated currents to those in cells with lipid-free pipette solution.

**Figure 2.**
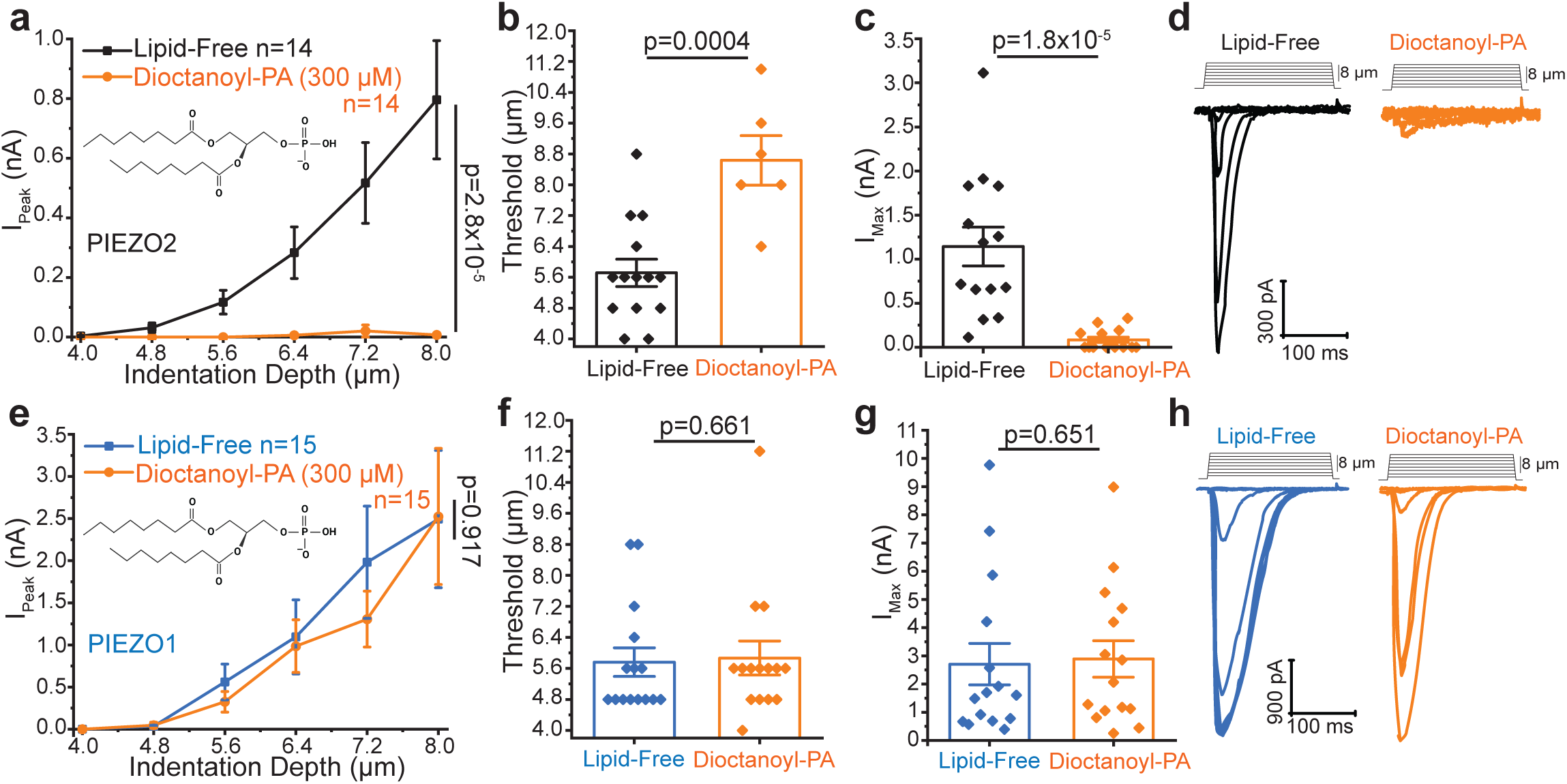
Phosphatidic acid inhibits PIEZO2 but not PIEZO1. Whole-cell patch-clamp experiments at -60 mV in N2A-Pz1-KO cells transiently transfected with PIEZO2-GFP or PIEZO1-GFP with patch-pipette solution supplemented with the indicated lipids as described in the methods section. **(a-d)** PIEZO2-GFP transfected cells supplemented with 300 μM dioctanoyl-phosphatidic acid (PA). **(a)** Current amplitudes are plotted (mean ± SEM) and statistical difference for the area under curve (AUC) for 4.0-8.0 µm stimuli calculated with the Mann-Whitney test. **(b)** Membrane indentation depth threshold to elicit mechanically-activated current (t-test). **(c)** Maximum current amplitudes (Mann-Whitney). **(d)** Representative current traces. **(e-h)** PIEZO1-GFP transfected cells supplemented with 300 μM dioctanoyl-phosphatidic acid. **(e)** Current amplitudes are plotted (mean ± SEM) and statistical difference of AUC for 4.0-8.0 µm stimuli calculated with the Mann-Whitney test. **(f)** Membrane indentation depth threshold to elicit mechanically-activated current (Mann-Whitney). **(g)** Maximum current amplitudes (Mann-Whitney). **(h)** Representative current traces. All bar graphs plotted with scatter and mean ± SEM.

First, we tested the effect of phosphatidic acid (**Fig. 1a**), which showed the most robust increase in TMEM120A transfected cells. Mechanically-activated PIEZO2 currents were robustly inhibited in experiments where we included the water-soluble short acyl chain dioctanoyl phosphatidic acid (300 μM) in the whole cell patch pipette, compared to control cells (**Fig. 2a-d**). In these experiments, we increased the stimulation depth in each cell until the seal was lost, but only plotted currents up to 8 μm in **Fig. 2a** for visual clarity. In the control group, 100% of cells displayed mechanically-activated currents while in the phosphatidic acid group only 42.9% of the cells responded. The mechanical threshold to evoke currents was significantly lower in control cells than in cells with intracellular phosphatidic acid application (**Fig. 2b**), and the maximal current, evoked by largest stimulus before seal rupture, was also significantly higher in control cells compared to phosphatidic acid treated cells (**Fig. 2c**). As shown in **Fig. 2e-h**, PIEZO1 currents were not affected by dioctanoyl phosphatidic acid. Both mechanical threshold (**Fig. 2f**), and maximal PIEZO1 currents (**Fig. 2g**) were similar in control and in phosphatidic acid treated cells. The lack of inhibition by phosphatidic acid is consistent with PIEZO1 channels also not being inhibited by TMEM120A^8^. The inactivation time constant (tau) was not different between control and phosphatidic acid treated cells for PIEZO2 (**Supplementary Fig. 3a**) and PIEZO1 (**Supplementary Fig. 3b**).

Next, we tested the effect of LPA, a lipid that also showed a significant increase in TMEM120A expression cells **(Fig. 1a)**. Inclusion of palmitoyl LPA (30 μM) in the patch pipette also inhibited PIEZO2 channels **(Fig. 3a-d)**. The area under the curve between 4 and 8 μm indentations was significantly lower in LPA treated cells (p=0.008) (**Fig. 3a**), and the mechanical threshold to evoke currents was significantly lower (p=0.001) in control cells than in cells with intracellular LPA application **(Fig. 3b)**. The maximal current evoked was also higher in control cells compared to LPA treated cells, but this difference was not statistically significant (p=0.08) (**Fig. 3c**). Intracellular application of LPA did not inhibit mechanically-activated PIEZO1 currents (**Fig. 3e-h**). Neither the threshold for mechanically-activated currents (**Fig. 3f**), nor the maximal PIEZO1 current (**Fig. 3g**) were different between control and LPA-treated cells. The inactivation time constant (tau) was not different between control and LPA treated cells for PIEZO2 (**Supplementary Fig. 3c**) and PIEZO1 (**Supplementary Fig. 3d**).

**Figure 3.**
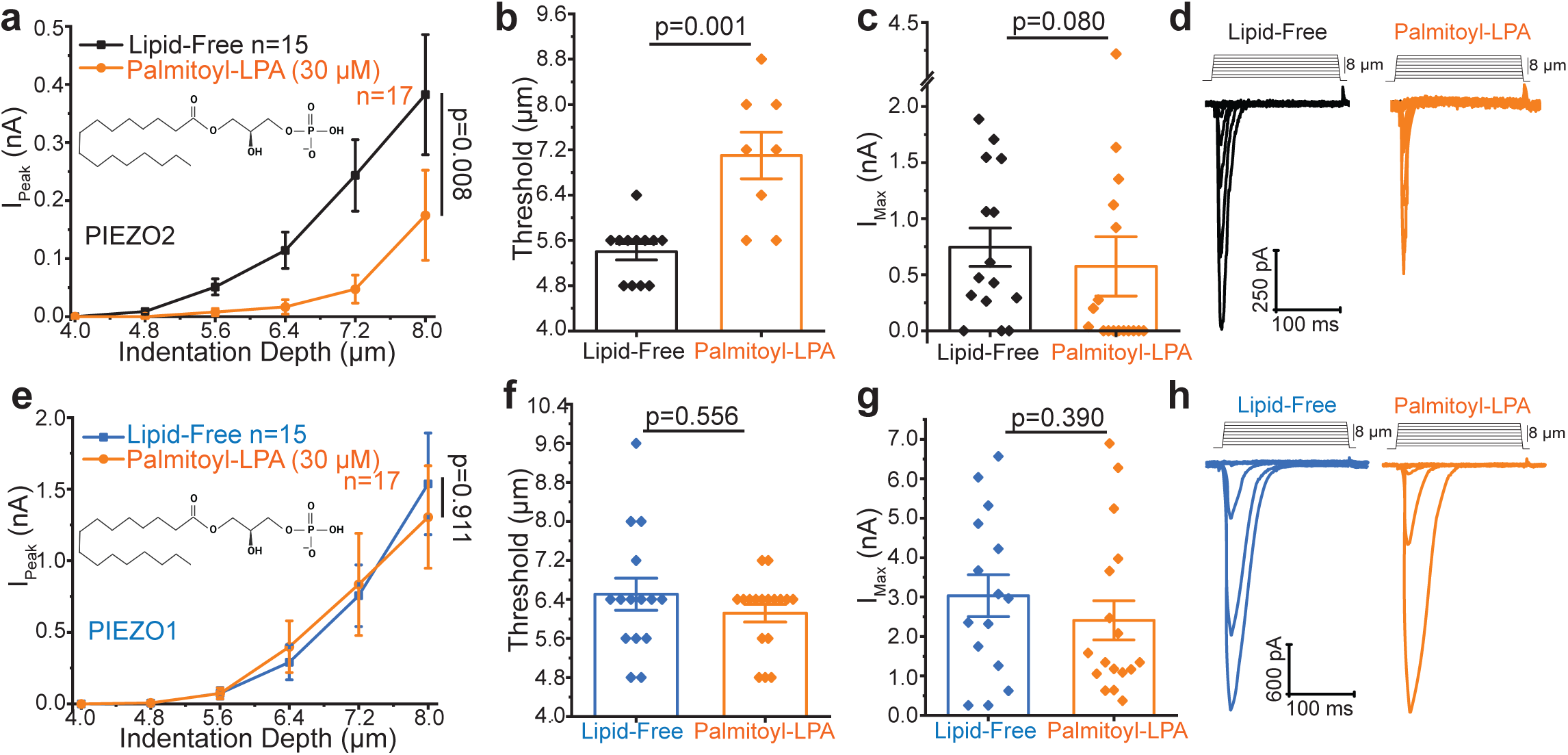
LPA inhibits PIEZO2 but not PIEZO1. Whole-cell patch-clamp experiments at -60 mV in N2A-Pz1-KO cells transiently transfected with PIEZO2-GFP or PIEZO1-GFP with patch-pipette solution supplemented with the indicated lipids as described in the methods section. **(a-d)** PIEZO2-GFP expressing cells supplemented with 30 μM palmitoyl-LPA. **(a)** Current amplitudes are plotted (mean ± SEM) and statistical difference of AUC for 4.0-8.0 µm stimuli calculated with the Mann-Whitney test. **(b)** Membrane indentation depth threshold to elicit mechanically-activated currents (Mann-Whitney). **(c)** Maximum current amplitudes (Mann-Whitney). **(d)** Representative current traces. **(e-h)** PIEZO1-GFP expressing cells supplemented with 30 μM palmitoyl-LPA. **(e)** Current amplitudes are plotted (mean ± SEM) and statistical difference of AUC for 4.0-8.0 µm stimuli calculated with the Mann-Whitney test. **(f)** Membrane indentation depth threshold to elicit mechanically-activated current (Mann-Whitney). **(g)** Maximum current amplitudes (Mann-Whitney). **(h)** Representative current traces. All bar graphs plotted with scatter and mean ± SEM.

Intriguingly, the lipids that showed the largest increase in TMEM120A transfected cells, were generated by enzymes that require acyl-CoA (**Fig. 1a**). This may suggest that TMEM120A provides acyl-CoA to these enzymatic steps, which would be consistent with ELOVL7 being an acyl-CoA elongase^20^. Long-acyl-chain CoA is known to modulate ion channels^21,22^ therefore we tested if including palmitoyl CoA (30 μM) in the whole cell patch pipette affects PIEZO2 activity. **Supplementary Fig. 3e-h**. shows that amplitudes of mechanically-induced currents in PIEZO2 expressing cells were similar to those in cells that were dialyzed with palmitoyl-CoA and control cells. The inactivation time constant (tau) for PIEZO2 currents was also not different between control and palmitoyl-CoA treated cells (**Supplementary Fig. 3i**). These data show that acyl-CoA is unlikely to be the lipid responsible for PIEZO2 inhibition upon TMEM120A expression.

### A non-metabolyzable analogue of phosphatidic acid and LPA inhibits PIEZO2 channels

Our experiments so far showed that phosphatidic acid and LPA inhibit PIEZO2 currents when delivered intracellularly. Phosphatidic acid and LPA are both charged, thus unlikely to rapidly cross the plasma membrane, and also metabolically unstable, and can be converted to other lipids inside and outside the cell. LPA also activates cell surface receptors, which would complicate interpretation of results with extracellular application. Carbacyclic Phosphatidic Acid (ccPA), is a metabolically stable analogue of phosphatidic acid and LPA that does not activate LPA receptors^23^. It was also reported that ccPA administration reduced mechanical allodynia in rats but whether ccPA affects PIEZO2 channel activity has not been tested^24^.To explore the possibility that extracellular application of this phosphatidic acid and LPA analogue can be used to modulate PIEZO2 activity, we tested the effect of long term (overnight) incubation with ccPA on mechanically-activated PIEZO2 currents. We incubated N2A-Pz1-KO cells expressing PIEZO2 with 300 μM ccPA, and we found that mechanically-activated currents were inhibited compared to control cells (**Fig. 4a-d**). Both maximal currents (**Fig. 4c**) and the area under the curve between 4 and 8 μm stimulations (**Fig. 4a**) were significantly lower in ccPA treated cells, but the threshold for mechanically-induced currents was not different (**Fig. 4b**).

**Figure 4.**
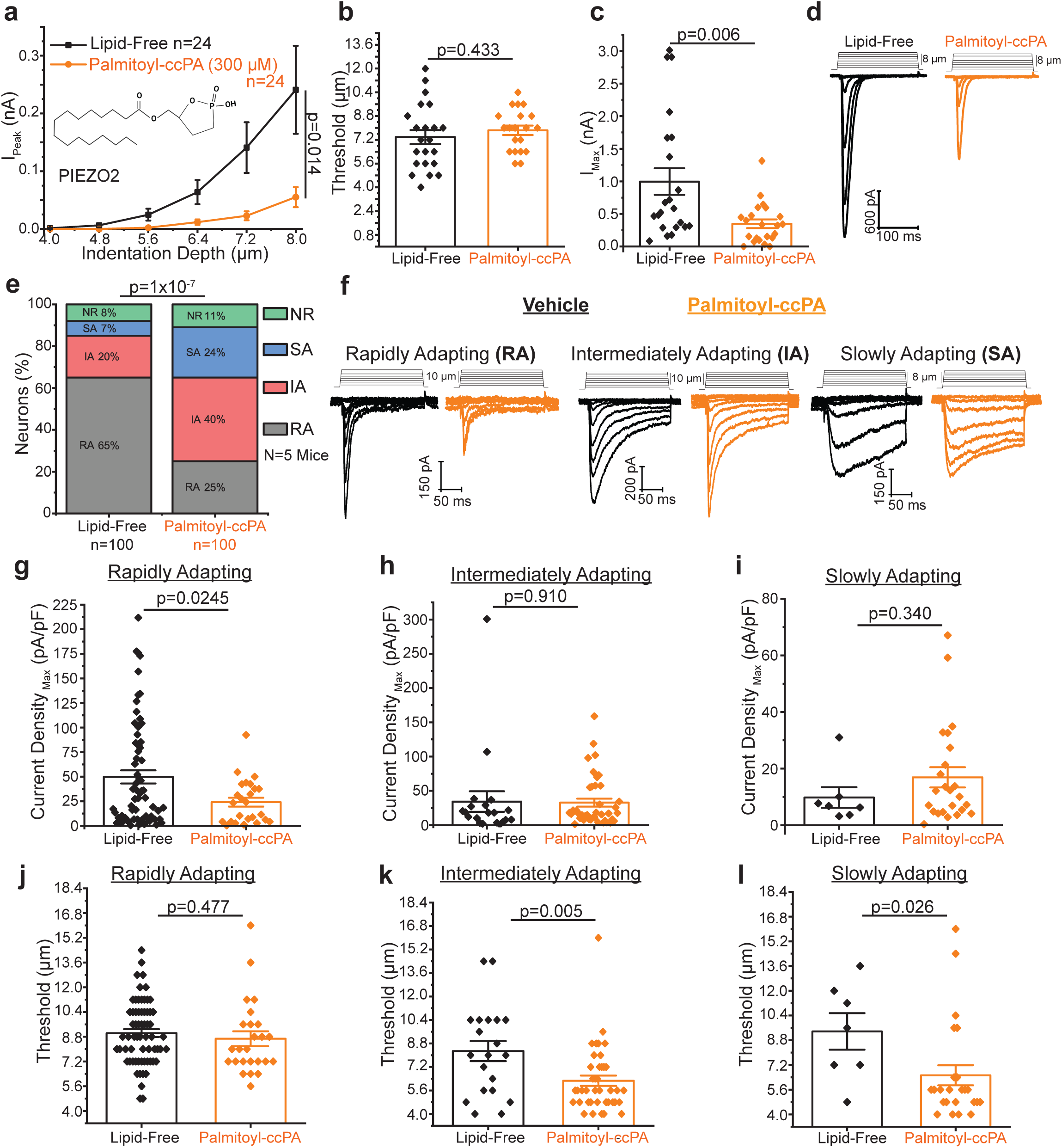
Extracellular application of ccPA inhibits PIEZO2 currents. Whole-cell patch-clamp experiments performed at -60 mV as described in the methods section. **(a-d)** N2A-Pz1-KO cells transiently transfected with PIEZO2-GFP mock treated (black) or treated with extracellular palmitoyl-ccPA overnight (orange). **(a)** Current amplitudes are plotted (mean ± SEM) and statistical difference the area under curve (AUC) for 4.0-8.0 µm stimuli calculated with the Mann-Whitney test. **(b)** Membrane indentation depth threshold to elicit mechanically-activated currents (t-test). **(c)** Maximum current amplitudes (Mann-Whitney). **(d)** Representative current traces. **(e-l).** Isolated mouse DRG neurons mock treated (black) or treated with extracellular palmitoyl-ccPA overnight (orange). **(e)** Proportion of rapidly adapting (RA), intermediately adapting (IA), slowly adapting (SA) and non-responders (NR). Chi-squared test. **(f)** Representative current traces. **(g)** Rapidly adapting maximum current density (t-test). **(h)** Intermediate adapting maximum current density (t-test). **(i)** Slowly adapting maximum current density (Mann-Whitney). **(j)** Membrane indentation depth threshold to elicit rapidly adapting current (t-test). **(k)** Membrane indentation depth threshold to elicit intermediately adapting current (t-test). **(l)** Membrane indentation depth threshold to elicit slowly adapting current (Mann-Whitney). All bar graphs plotted with scatter and mean ± SEM.

Next, we tested if endogenous rapidly adapting mechanically-activated currents in DRG neurons are inhibited by ccPA. **Fig. 4e** shows that the proportion of neurons displaying rapidly adapting MA currents decreased from 65% to 25 % when the cells were treated overnight with ccPA. In the neurons that displayed rapidly adapting mechanically-activated currents, the maximal current density was significantly lower in cells treated with ccPA (**Fig. 4g**). There was a compensatory increase in the proportion of non-responding cells in the ccPA treated group. The proportion of slowly adapting and intermediate currents also increased (**Fig. 4e**), but their amplitudes were not different in ccPA treated and control cells (**Fig. 4h,i**). The mechanical threshold of rapidly adapting mechanically-activated currents did not change in ccPA treated cells (**Fig. 4j**), but the mechanical threshold of intermediate and slowly adapting currents significantly decreased (**Fig. 4k,l**). Since ccPA, PA and LPA treatment did not change the inactivation time constant of expressed recombinant PIEZO2 currents (**Supplementary Fig. 4a**), the increase in the proportion of intermediate and slowly adapting current is likely due to unmasking of slowly adapting and intermediate currents by PIEZO2 inhibition, and potentially a compensatory increase in those currents. Overall, these data indicate that ccPA treatment inhibits both recombinant and endogenous PIEZO2-mediated mechanically-activated currents.

### Phospholipase D (PLD) regulates PIEZO2 but not PIEZO1 channels

Phosphatidic acid can also be generated by phospholipase D enzymes from phosphatidylcholine^25^. To test if PLD activation modulates PIEZO2 activity, we used the opto-PLD system, where PLD activity in the plasma membrane can be induced by blue light^26^. This system is based on recruitment of a PLD enzyme fused to the CRY2 plant protein which undergoes heterodimerization with CIBN in response to blue light. CIBN is targeted to the plasma membrane, and therefore blue light induces a translocation of PLD to the plasma membrane (**Fig. 5a**). We found that PIEZO2 activity is inhibited by blue light exposure in cells expressing the two components of the opto-PLD system (**Fig. 5b,c,e**). When the components of the Opto-PLD system containing a catalytically inactive PLD mutant (H170A) were co-transfected, blue light illumination did not inhibit PIEZO2 (**Fig. 5b,c,d**). Mechanically-activated PIEZO1 currents was not inhibited by activating the opto-PLD system with blue light (**Supplementary Fig. 5**).

**Figure 5.**
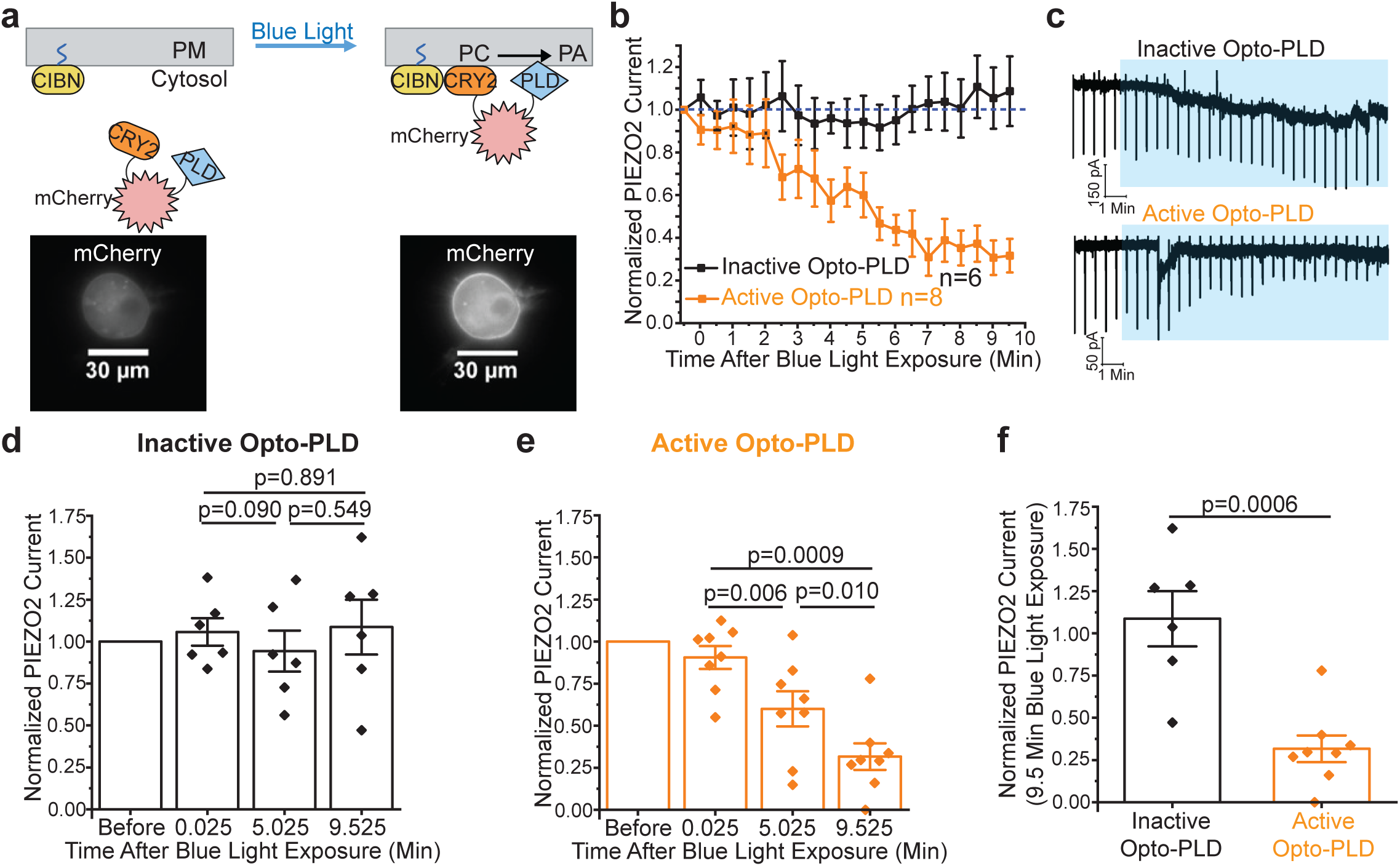
Optogenetic activation of PLD inhibits PIEZO2. Whole-cell patch-clamp experiments at -60 mV in N2A-Pz1-KO cells transfected with PIEZO2-GFP and active Opto-PLD (black) or inactive Opto-PLD (orange; H170A) as described in the methods section. **(a)** Scheme of blue light activation of the Opto-PLD system (generated with BioRender) with representative images for mCherry fluorescence (60x) before and after blue light exposure. **(b)** PIEZO2-GFP currents with fixed, continuous membrane indentations after blue light exposure normalized to currents before blue light. **(c)** Representative current traces. Blue shaded area indicates blue light exposure. - Downward deflections indicate mechanically-activate Piezo2 currents **(d)** Scatter and mean ± SEM for PIEZO2-GFP co-expressed with inactive Opto-PLD after blue light exposure (Repeated-Measures ANOVA, F=0.365, df=2,10, p=0.702; paired t-tests displayed). **(e)** Scatter and mean ± SEM for PIEZO2-GFP co-expressed with active Opto-PLD after blue light exposure (Repeated-Measures ANOVA, F=21.386, df=2,14, p=5.546x10^-5^; paired t-tests displayed). **(f)** Scatter and mean ± SEM of PIEZO2-GFP with inactive or active Opto-PLD after 9.5 min of blue light exposure (t-test).

Our data so far show that acutely activating PLD inhibits PIEZO2 activity. Next, we tested if endogenous basal PLD activity contributes to regulation of PIEZO2 channels. To achieve this, we treated PIEZO2 expressing N2A-Pz1-KO cells with the broad spectrum PLD inhibitor FIPI (500 nM). As shown in **Figure 6b-e**, mechanically-activated PIEZO2 currents were significantly increased by FIPI treatment. The mechanical threshold to evoke PIEZO2 currents was significantly lower in FIPI treated cells (**Fig. 6c**) and the maximal current amplitude was significantly higher (**Fig. 6d**).

**Figure 6.**
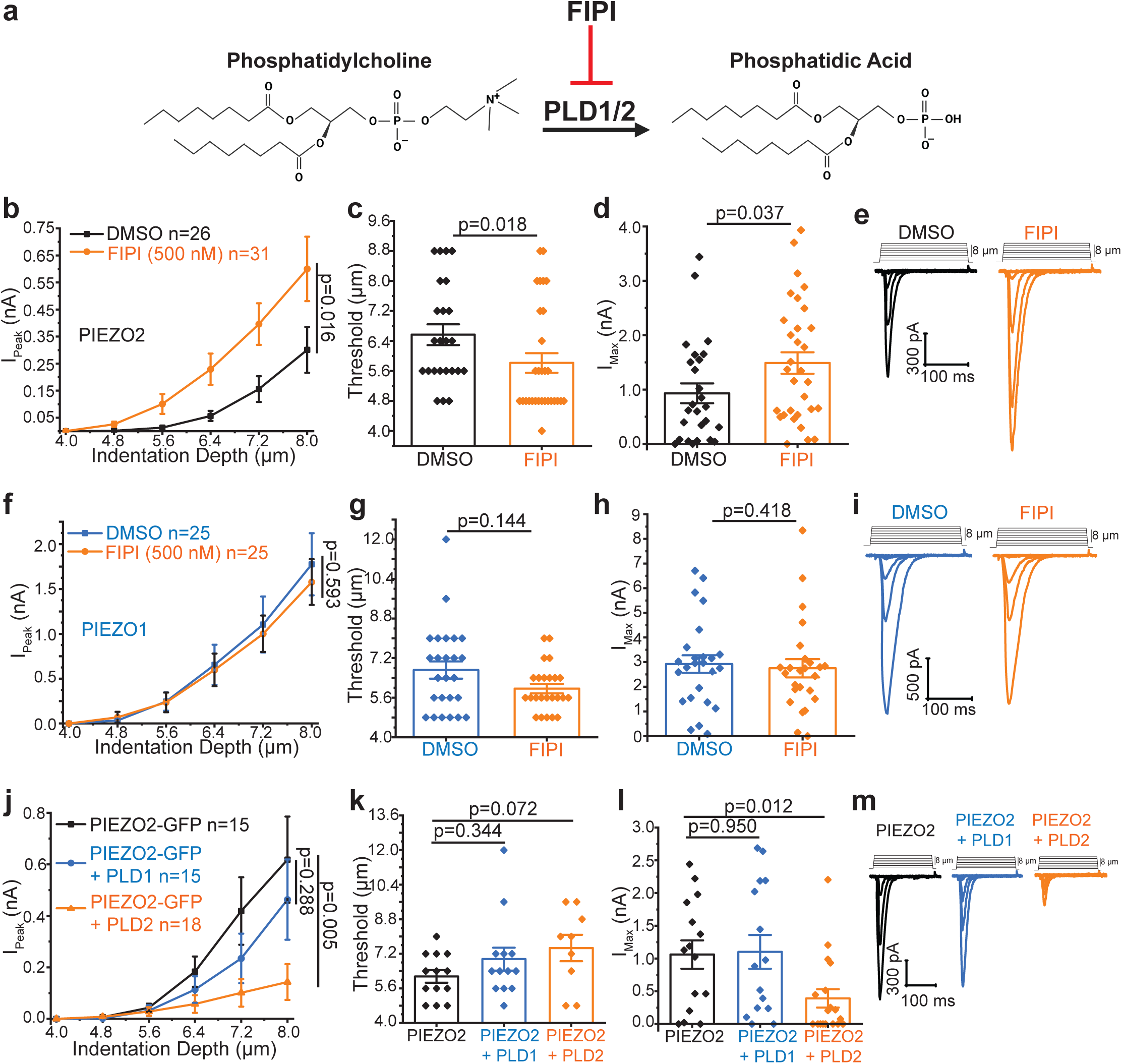
Phospholipase D (PLD) negatively regulates PIEZO2 but not PIEZO1. Whole-cell patch-clamp experiments at -60 mV in transiently transfected N2A-Pz1-KO cells as described in the methods section. **(a)** Scheme of PLD inhibition by FIPI. **(b-e)** N2A-Pz1-KO cells transiently transfected with PIEZO2-GFP and were treated with DMSO (black) or 500 nM FIPI (orange) for 30 min, see methods for details. **(b)** Current amplitudes are plotted (mean ± SEM) and statistical difference of AUC for 4.0-8.0 µm stimuli calculated with the Mann-Whitney test. **(d)** Membrane indentation depth threshold to elicit mechanically-activated current (Mann-Whitney). **(d)** Maximum current amplitudes (Mann-Whitney). **(e)** Representative current traces. **(f-i)** N2A-Pz1-KO cells transiently transfected with PIEZO1-GFP treated with 500 nM FIPI (orange) for 30 min or the vehicle DMSO (blue). **(f)** Current amplitudes are plotted (mean ± SEM) and statistical difference of AUC for 4.0-8.0 µm stimuli calculated with the Mann-Whitney test. **(g)** Membrane indentation depth threshold to elicit mechanically-activated currents (Mann-Whitney). **(h)** Maximum current amplitudes (Mann-Whitney). **(i)** Representative current traces. **(j-m)** N2A-Pz1-KO cells transiently transfected with PIEZO2-GFP alone (black), or together with PLD1 (blue), or PLD2 (orange). **(j)** Current amplitudes are plotted (mean ± SEM) and statistical difference of AUC for 4.0-8.0 µm stimuli calculated was with the Mann-Whitney test. **(k)** Membrane indentation depth threshold to elicit mechanically-activated current (Kruskal-Wallis, χ^2^=3.298, df=2, p=0.0192; Mann-Whitney tests displayed). **(l)** Maximum current amplitudes (Kruskal-Wallis, χ^2^=7,906, df=2, p=0.0192; Mann-Whitney tests displayed). **(m)** Representative current traces. All bar graphs plotted with scatter and mean ± SEM.

PIEZO1 currents on the other hand were not potentiated by FIPI (**Fig. 6f-i**), which is consistent with our finding that PIEZO1 currents were not affected by phosphatidic acid **(Fig. 2e)**. The inactivation time constant (tau) was similar in FIPI treated and control cells for both Piezo2 and Piezo1 currents (**Supplementary Fig. 6 a,b**).

There are two mammalian PLD enzymes, PLD1 and PLD2. To test which isoform is responsible for the basal regulation of PIEZO2 activity, we coexpressed PLD1 or PLD2 with PIEZO2 in N2A-Pz1-KO cells. **Figure 6j-m** shows that coexpression of PLD2, but not PLD1 significantly inhibited PIEZO2 activity. The maximal current in PLD2 expressing cells was significantly lower than in control cells (**Fig. 6l**). Mechanical threshold was also higher in PLD2 expressing cells (**Fig. 6k**), but it did not reach statistical significance (p=0.07). The inactivation time constant (tau) was similar in cells expressing PLD1 or PLD2 and in control cells (**Supplementary Fig. 6c**). The inhibition of PIEZO2 currents by PLD2 but not by PLD1 is consistent with PLD2, but not PLD1 showing constitutive activity^27^.

### PLD modulates native rapidly adapting mechanically-activated currents in peripheral sensory neurons and mechanical sensitivity in mice

Our data with recombinant PIEZO2 currents show that these channels can be modulated by the PLD pathway. Next, we tested if endogenous PLD activity also regulates PIEZO2 channels in their native environment. First, we performed experiments in human induced pluripotent stem cell (hiPSC)-derived sensory neurons (Anatomic Incorporated)^28^. Consistent with earlier results^28^, these cells displayed predominantly rapidly adapting mechanically-activated currents. As shown in **Figure 7a**, mechanically-activated current amplitudes were significantly higher in FIPI-treated cells, compared to control cells. Mechanical threshold was significantly lower (**Fig. 7b**), and maximal current density was significantly higher (**Fig. 7c**) in FIPI-treated cells compared to control cells.

**Figure 7.**
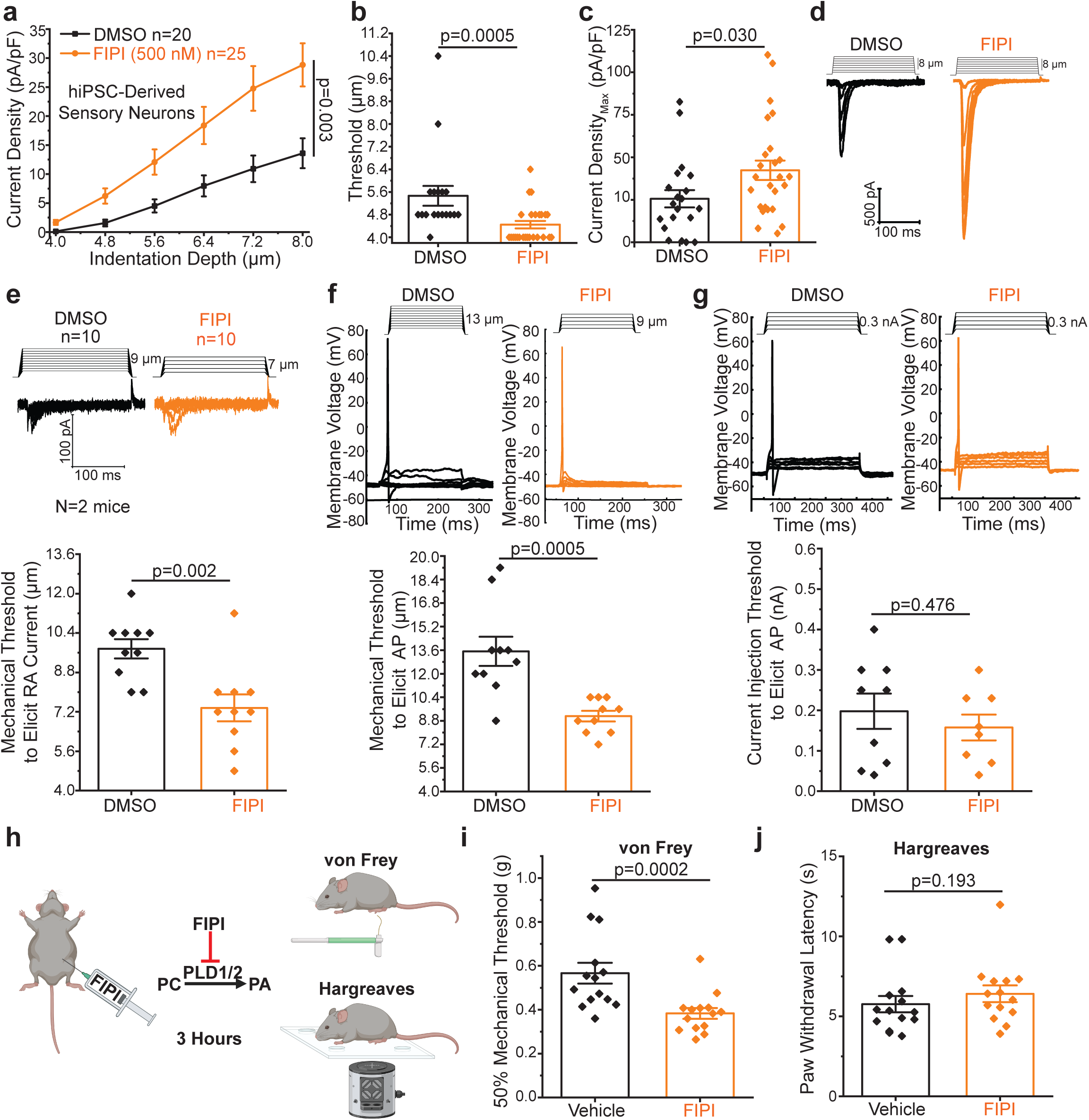
PLD modulates native rapidly adapting mechanically-activated currents in peripheral sensory neurons. **(a-d)** Whole-cell patch-clamp at -60 mV in hiPSC-sensory neurons treated with 500 nM FIPI (orange) or DMSO (black) as described in the methods section for details. **(a)** Current densities are plotted (mean ± SEM) and statistical difference of AUC for 4.0-8.0 µm stimuli calculated with the Mann-Whitney test. **(b)** Membrane indentation depth thresholds to elicit mechanically-activated current (Mann-Whitney). **(c)** Maximum current densities (Mann-Whitney). **(d)** Representative current traces. **(e)** Whole-cell patch-clamp at -60 mV in isolated mouse DRG neurons treated with DMSO (black) or FIPI (orange) as described in the methods section. Representative current traces (top), and membrane indentation depth thresholds to elicit mechanically-activated RA currents (bottom) (t-test). **(f-g)** Whole-cell current-clamp in isolated mouse DRG neurons treated with DMSO (black) or FIPI (orange) as described in the methods section. **(f)** Representative traces (top), and membrane indentation depth threshold to elicit an action potential (bottom) (t-test). **(g)** Representative traces (top), and current injection threshold to elicit an action potential (bottom) (t-test) in the same cells used in panels E and F. In three cells the seal was lost before current injections could be performed All bar graphs plotted with scatter and mean ± SEM. **(h-j)** Mechanical testing using Von Frey filaments and thermal testing using the Hargreaves apparatus were performed on mice receiving an I.P. injection of FIPI (n=14) or vehicle (n=14) as described in the methods section. **(h)** Experimental design (generated with BioRender). **(i)** von Frey testing 50% mechanical threshold (Mann-Whitney). **(j)** Hargreaves paw withdrawal latency (Mann-Whitney).

Next, we tested the effect of the PLD inhibitor FIPI in DRG neurons displaying rapidly adapting mechanically-activated currents. We first performed whole-cell voltage clamp experiments to identify neurons that display rapidly adapting mechanically-activated currents, using the same increasing indentation depth protocol we used in our experiments so far, but we stopped mechanical stimulation shortly after clear rapidly adapting currents appeared, to avoid losing the seal. The mechanical threshold to elicit rapidly adapting currents was significantly lower in FIPI-treated cells compared to control cells (**Fig. 7e**). Then on the same neurons we performed current clamp measurements to test mechanical threshold to elicit action potentials. As shown in **Figure 7f**, the indentation depth threshold to elicit action potentials was significantly lower in FIPI-treated cells, than in control neurons. We also tested the current injection threshold to elicit action potentials. As shown in **Figure 7g**, the current injection thresholds to elicit action potentials were similar in FIPI-treated and control cells. The resting membrane potential was also not significantly different between control (of -50.4 ± 0.84 mV) and FIPI treated neurons (-48.2 ± 1.56, p=0.252) (**Supplementary Fig. 7c**). These data indicate that the effect of FIPI on the mechanical threshold was not due to the increased general excitability of DRG neurons.

To test if PLD inhibition by FIPI can modulate PIEZO2-mediated touch sensation, we injected mice with FIPI intraperitoneally, and tested their mechanical sensitivity using von Frey filaments (**Fig. 7h**). We found that mice injected with FIPI had significantly lower 50% mechanical threshold to withdraw their hind paw from von Frey filaments (**Fig. 7i**). This is consistent with FIPI potentiating PIEZO2 currents and lowering firing threshold evoked by mechanical stimuli in DRG neurons. Paw withdrawal latency to thermal stimuli on the other hand were not significantly different in FIPI-injected and control mice (**Fig. 7j**), indicating that the effect of FIPI was specific to mechanical stimuli.

Overall, these data indicate that basal PLD activity modulates PIEZO2 channels in DRG neurons, and mechanical sensitivity *in vivo*.

## DISCUSSION

PIEZO1 and PIEZO2 are mechanically-activated ion channels and they play roles in a wide variety of physiological functions. Despite their biomedical importance, no endogenous or exogenous small molecule inhibitor has been identified so far that selectively inhibits PIEZO2 over PIEZO1. Here we build on our earlier work showing that co-expression of the TMEM120A protein inhibits PIEZO2, but not PIEZO1, and we identify phosphatidic acid and LPA as endogenous inhibitors of PIEZO2. We also find that phosphatidic acid generated by PLD enzymes inhibits PIEZO2, but not PIEZO1, thus we identify the PLD pathway as a regulator of PIEZO2 activity.

PIEZO1 and PIEZO2 share a similar structure of a propeller-shaped trimer, with 38 transmembrane domains in each subunit^29^. The membrane footprint of PIEZO proteins is larger than that of any other known ion channel, implying that membrane lipids could greatly influence their behavior. However, research on the impact of membrane lipids on PIEZO channel function is still in early stages. Various dietary fatty acids have been shown to modulate activity of both PIEZO1 and PIEZO2 channels, but none of those lipids showed selectivity for PIEZO2 over PIEZO1. The dietary saturated fatty acid margaric acid (C17:0) has been shown to inhibit both PIEZO1 and PIEZO2 activity, with a higher potency on PIEZO1^17,30^. Linoleic acid (C18:2), an unsaturated dietary fatty acid on the other hand was shown to increase the activity of both PIEZO1 and PIEZO2^28,30^.

The polyunsaturated fatty acid eicosapentanoic acid (EPA, 20:5) decreased the inactivation time constant for wild-type and gain of function mutant PIEZO1^30^ and PIEZO2 channels^31^. The sphingomyelinase enzyme has been reported to alter PIEZO1’s inactivation kinetics in vascular endothelial cells by producing ceramides^32^.

Recent computational studies using coarse-grained simulation of PIEZO1 and PIEZO2 proteins in a lipid membrane similar in composition to mammalian plasma membranes found that negatively charged membrane lipids are enriched in the vicinity of the propeller shaped arms of the PIEZO proteins^33–35^. These lipids include phosphatidic acid, phosphatidylserine, phosphatidylinositol and phosphatidylinositol phosphates.

Other studies on PIEZO1 also concluded that phosphatidylinositol 4,5-bisphosphate [PI(4,5)P_2_] is enriched in the vicinity of the PIEZO arms^36,37^, which is not surprising, given the large number of positive charged residues in the cytoplasm inner membrane interface of both channels. Depletion of PI(4,5)P_2_ Ca^2+^-induced activation of phospholipase C (PLC) enzymes upon TRPV1 channel activation have been shown to inhibit PIEZO1 and PIEZO2^38^. Dephosphorylation of PI(3,5)P_2_ by the phosphoinositide phosphatase myotubularin related protein-2 was also reported to reduce PIEZO2 activity^39^.

All computational studies found multiple potential binding sites for negatively charged lipids, with most sites binding several lipids with varying predicted affinity and similar, yet not identical sites among the different studies. The potential complexity of lipid interaction with PIEZO2 makes it hard to pinpoint potential binding sites on the channel, and further studies will be required to identify lipid binding sites responsible for the effect of phosphatidic acid and LPA.

We identified phosphatidic acid and LPA as inhibitors of PIEZO2 by dissecting the mechanism of inhibition by TMEM120A. TMEM120A (a.k.a.TACAN) was proposed to be a component of slowly adapting mechanically-activated ion channels responsible for mechanical pain^9^. This proposal was based on the appearance of mechanically-activated currents in cells transfected with TMEM120A, and on the finding that siRNA knockdown of TMEM120A reduced the proportion of slowly adapting mechanically-activated currents in DRG neurons. These findings however were not reproduced by other laboratories^8,10–13,40^, raising doubts about TMEM120A functioning as a mechanically-activated channel. Structural studies^10,12,13,40,41^ indicated that TMEM120A shares structural homology to the long chain fatty acid elongase enzyme ELOVL7^14^.

Other studies indicated that TMEM120A is involved in adipocyte differentiation^42^ and triglyceride production^42^, and its genetic deletion from adipocytes resulted in lipodystrophy in mice^43^. These studies indicated that TMEM120A is located in the nuclear envelope, and suggested that its effect on adipocytes was mediated by affecting expression of other genes, see our recent review on TMEM120A for further discussion^15^. Our data show that TMEM120A overexpression increases intermediates in the Kennedy pathway of triglyceride synthesis. It will require further studies if this effect is due to an enzymatic activity of TMEM120A, or modifying expression of other genes.

Our primary goal here was to take advantage of the selective inhibition of PIEZO2 by TMEM120A, and identify potential channel inhibitor molecules. Thus we focused on the two lipids that showed the most robust increase in TMEM120A expressing cells, phosphatidic acid and LPA, and found that intracellular application of either of these lipids inhibited PIEZO2, but not PIEZO1.

Phosphatidic acid, other than being an intermediate of triglyceride synthesis in the Kennedy pathway, can also be generated by phospholipase D (PLD) enzymes that cleave the head group from phospholipids, mainly phosphatidylcholine, leaving the phosphate on the glycerol backbone and thus generating phosphatidic acid.

Phosphatidic acid has been thoroughly studied in plant hormonal signaling^44^, and has been implicated in modulating cellular processes such as membrane traffic, endocytosis and exocytosis in mammals, but its precise role in these process is not well defined^45^.

There have also been sporadic reports on phosphatidic acid affecting ion channels^46,47^, but this lipid has been understudied in mammalian signaling compared to phosphoinositides and DAG. Specifically, PLD2 has been implicated in the positive regulation of mechanosensitive TREK-1 channels^47^. Given that TREK-1 is a mechanically-activated hyperpolarizing channel, which is positively regulated by PLD2, one may argue that TREK channel modulation contributes to the effects of FIPI in modulating mechanosensitivity in current clamp and behavioral experiments. Our findings that neither resting membrane potential, nor current injection threshold was different in FIPI-treated and control neurons, however, argue against the involvement of TREK-1 in the effects of FIPI on mechanically-induced electrical activity and mechanosensitivity in behavioral experiments.

LPA, besides being an intermediate in the Kennedy pathway of triglyceride synthesis, is also well known as an extracellular signaling molecule, acting on cell surface receptors. Extracellular LPA is generated by either PLA1/2 enzymes hydrolyzing extracellularly facing PA, or by conversion of lysophosphatidyl choline (also produced by PLA1/2) to LPA by autotaxin enzymes^48^. Extracellular LPA is thought to be generated by these enzymatic reactions on the extracellular surface of the plasma membrane^48^, and it is unlikely that LPA generated inside the cell crosses the plasma membrane in sufficient quantities to serve as an extracellular signal^49^. LPA has also been reported to activate TRPV1 and TRPA channels via direct binding^50,51^.

PLD enzymes generate phosphatidic acid by removing the head group of phosphatidylcholine **(Fig. 5a)**. There are two PLD enzymes in mammals. PLD2 is thought to have constitutive activity, while PLD1 is tightly regulated by a variety of factors including the small G-proteins Arf and Rho, as well as Protein Kinas C (PKC)^27^. PLD-s are involved in a variety of cellular functions, including cell metabolism, migration, and exocytosis^27^. Even though both PLD-s are expressed in DRG neurons^52-^ _54_, the role of PLD enzymes in somatosensory physiology, and specifically in the regulation of PIEZO2 channels, is completely unexplored. Here we show that optogenetic activation of PLD inhibits PIEZO2, but not PIEZO1, demonstrating that in principle, PLD activity can influence PIEZO2 channel activity. We also find that the PLD inhibitor FIPI increases mechanically-activated PIEZO2, but not PIEZO1 currents, indicating that basal PLD activity modulates PIEZO2 activity. PLD inhibition by FIPI also increased the amplitudes of endogenous rapidly adapting mechanically-activated currents in hiPSC-derived peripheral sensory neurons, and reduced the mechanical threshold for action potential firing in mouse DRG neurons that had rapidly adapting mechanically-activated currents. Consistent with these data FIPI injection decreased the mechanical threshold in experiments using von Frey filaments, a behavioral assay that was shown to be dependent on PIEZO2 channels^55^.

In conclusion, we show that phosphatidic acid and LPA inhibits PIEZO2, but not PIEZO1 activity, and identify the PLD pathway as a modulator of PIEZO2 channels and mechanical sensitivity in mice. The specific inhibition of PIEZO2 over PIEZO1 by phosphatidic acid and LPA may serve as a basis for future work to identify clinically useful specific PIEZO2 inhibitors.

## METHODS

### Animals

All animal procedures were approved by the Institutional Animal Care and Use Committee at Rutgers New Jersey Medical School. Mice were kept in a barrier facility under a 12/12 hour light/dark cycle. Adult (8-12 weeks old) wild-type C57BL/6 mice (The Jackson Laboratory) of both sexes were used for both experiments.

### Cell lines

Neuro2A-*Piezo1*-KO (N2A-Pz1-KO) cells in which endogenous *Piezo1* was deleted using CRISPR were a kind gift from Drs. Valeria Vasquez and Gary Lewin. These cells were cultured in Dulbecco’s Modified Eagle Medium/F-12 (HAM) supplemented with 1% penicillin-streptomycin and 10% FBS at 37°C with 5% CO_2_. These cells were transiently transfected plasmids as described below using either Effectene (Qiagen, Cat. Nu. 301425) or PEI-Max (Polysciences Inc, Cat. Nu. 24765-100). For whole-cell electrophysiology and TIRF experiments, these cells were split onto poly-L-lysine coated glass coverslips 24 hours after transfection and used for experiments 48 total hours after transfection. hiPSC derived peripheral sensory neurons (RealDRG^TM^) were differentiated and cultured according to manufacturer’s (Anatomic Incorporated) instructions (briefly described below) before whole-cell patch clamp experiments.

### Lipidomic LC-MS/MS Analysis

N2A-Pz1-KO cells were transfected using PEI-Max (Polysciences Inc, Cat. Nu. 24765-100) with 1500 ng ptdTomato-N1 vector, 1500 ng TMEM120A-Tom, or 1500 ng TMEM120B. Cells were harvested 48 hours after transfection. One million cells per group were centrifuged, and washed with ice-cold PBS. Each cell pellet was resuspended in 500 µL of resuspension buffer (0.05 M HCl, 49% Methanol, 1% SPLASH Lipidomic Mass Spec Standard [Avanti Polar Lipids, Cat. Nu. 330707]), and 1 mL methyl-tert-butyl-ether was added. Samples were vortexed for 30 seconds, and 600 µL of the organic layer was transferred to a new tube. Samples were allowed to dry overnight and stored at -80°C.

The reversed-phase separation was performed on a Vanquish Horizon UHPLC system (Thermo Fisher Scientific, Waltham, MA) with a Poroshell 120 EC-C18 column (150 mm × 2.1 mm, 2.7 μm particle size, Agilent InfinityLab, Santa Clara, CA) using a gradient of solvent A (90%:10% H2O:MeOH with 34.2 mM acetic acid, 1 mM ammonium acetate, pH 9.4), and solvent B (75%:25% IPA:methanol with 34.2 mM acetic acid, 1 mM ammonium acetate, pH 9.4). The gradient was 0 min, 25% B; 2 min, 25% B; 5.5 min, 65% B; 12.5 min, 100% B; 19.5 min, 100% B; 20.0 min, 25% B; 30 min, 25% B. The flow rate was 200 μl/min. The injection volume was 5 μL and the column temperature was 55 °C. The autosampler temperature was set to 4°C and the injection volume was 5µL. The full scan mass spectrometry analysis was performed on a Thermo Q Exactive PLUS with a HESI source which was set to a spray voltage of -2.7kV under negative mode and 3.5kV under positive mode. The sheath, auxiliary, and sweep gas flow rates of 40, 10, and 2 (arbitrary unit) respectively. The capillary temperature was set to 300°C and aux gas heater was 360°C. The S-lens RF level was 45. The m/z scan range was set to 100 to 1200 m/z under both positive and negative ionization modes. The AGC target was set to 1e6 and the maximum IT was 200 ms. The resolution was set to 140,000 at m/z 200. The lipidomic annotation and quantitation was performed using MS-DIAL^56,57^.

For the calculation of total LPA (0), PA (0), DG (0) and TG (0) in Figure 1, the total ion counts of 5 LPA, 9 PA, 8 DG and 13 TG species were summed. The same method is applied when calculating the total LPA (1), LPA (2-6), PA(1), PA(2-12), DG(1), DG(2-5), TG(1) and TG(2-18) levels for Supplementary Fig. 1.

### Whole-Cell Patch Clamp

Whole-cell patch-clamp recordings were performed at room temperature (22-24°C) as previously described^58^. Briefly, patch pipettes were prepared from borosilicate glass capillaries (Sutter Instrument, Cat. Nu. BF150-75-10) using a P-97 pipette puller (Sutter Instrument) and had a resistance of 2.0-7.0 MΩ. After forming GΩ-resistance seals, the whole-cell configuration was established. All electrophysiology recordings were performed with an Axopatch 200B amplifier (Molecular Devices) and pClamp 11.2.

Currents were filtered at 5 kHz using low-pass Bessel filter of the amplifier and digitized using a Digitata 1,440 unit (Molecular Devices). All voltage clamp measurements were performed at a holding voltage of -60 mV. For all current clamp measurements, cells that had a resting membrane potential more positive than -40 mV were excluded.

All measurements were performed using an extracellular solution containing 137 mM NaCl, 5 mM KCl, 1 mM MgCl_2_, 2 mM CaCl_2_, 10 mM HEPES, and 10 mM glucose (pH adjusted to 7.4 with NaOH). The patch pipette solution for voltage clamp (in heterologous systems) and current clamp measurements contained 140 mM K+ gluconate, 1 mM MgCl_2_, 2 mM Na_2_ATP, 5 mM EGTA, and 10 mM HEPES (pH adjusted to 7.3 with KOH). For mechanically-activated voltage clamp measurements in DRG neurons, the following cesium-based solution was used: 133 mM CsCl, 1 mM MgCl_2_, 1 mM CaCl_2_, 5 mM EGTA, 4 mM Na_2_ATP, 10 mM HEPES (pH adjusted to 7.3 with COH).

Mechanically-activated currents were measured from N2A-Pz1-KO cells transiently transfected with 500 ng / cDNA construct using Effectene (Qiagen, Cat. Nu. 301425), isolated mouse DRG neurons, and hiPSC-derived peripheral sensory neurons (Anatomic Incorporated, Cat. Nu. 1020F1-1M) as previously described^58^. Briefly, mechanical stimulation was performed using a heat-polished glass pipette (tip diameter about 3 µm) controlled by a piezoelectric crystal drive (Physik Instrumente) positioned at 60° to the surface of the cover glass. The probe was positioned so that 10-µm movement did not visibly contact the cell but an 11.5-µm stimulus produced an observable membrane deflection. Cells received a membrane indentation lasting 200 ms at a depth of 4.0 Δ +0.8 µm / 10 sec. For measurements using Opto-PLD (AddGene, Active Opto PLD Cat. Nu. 140114, Inactive Opto-PLD Cat Nu. 140061), an indentation depth that produced consistent and submaximal current was applied to the cells every 30 seconds. To reach an intensity of blue light sufficient to activate the Opto-PLD system, an LED light source (CoolLED, pE-300) and 60x oil objective (N.A.: 1.49) were used. Measurements from cells that showed significant swelling after repetitive mechanical stimulation or had substantially increased leak current were discarded.

For intracellular application of lipids, the patch pipette solution described above was supplemented with the denoted lipid. Once the whole-cell configuration was established, we waited 5 minutes for lipids to dialyze into the cell before measurements were taken. Patch pipette solution was used as a solvent for dioctanoyl-phosphatidic acid (Avanti Polar Lipids, Cat. Nu. 830842P), palmitoyl-LPA (Cayman Chemical, Cat. Nu. 10010094) and palmitoyl-CoA (MilliporeSigma, Cat. Nu. P9716) in these experiments. For extracellular application of lipids, DMEM/F-12 (HAM) was used as a solvent for palmitoyl-ccPA (Cayman Chemical, Cat Nu. 10010293) and cells were cultured for 24 hours in lipid-supplemented media before measurements were taken. For pharmacological inhibition of PLDs, cells were incubated in DMEM/F-12 (HAM) containing 500 nM FIPI (Cayman Chemical, Cat. Nu. 13563) or an equivalent volume of DMSO (0.005%) for 30 minutes and were patched in extracellular solution containing 500 nM FIPI. To inhibit GPAT, cells were incubated overnight with media supplemented with 100 µM FSG67 (Focus Biomolecules, Cat. Nu. 10-4577) or the equivalent volume of DMSO (0.1 %).

Data were collected and analyzed in Clampfit 11.2 software including the quantification of peak current amplitudes for each indentation depth. To compare the peak current amplitude data for the indentation depth range where the majority of cells survived (4.0-8.0 µm), AUC calculations were made in Origin 2021. The AUC for these indentation depths was calculated for each cell using the peak current amplitudes elicited through 4.0-8.0 µm indentation depths. The AUC values were compared between groups to evaluate any statistical differences for these stimulations.

The inactivation time constant (tau) for mechanically-activated currents were measured in Clampfit 11.2 software by fitting an exponential decay function. We used the currents evoked by the third stimulation after the threshold in most experiments. Except in the cells where only the two largest stimuli evoked a current, in which case we measured the tau using the largest stimulus (provided it reached 40 pA).

### DRG Neurons

DRG neurons were isolated as previously described^58^. Briefly, mice were anesthetized with an i.p. injection of ketamine (100 mg/kg) and xylazine (10 mg/kg) and perfused via the left ventricle with ice-cold Hank’s Buffered Salt Solution (HBSS). DRGs were harvested from all spinal segments after laminectomy and removal of the spinal column and maintained in ice-cold HBSS for the duration of the isolation. Isolated ganglia were cleaned from excess nerve tissue and incubated with type 1 collagenase (3 mg/mL) and dispase (5 mg/mL) in HBSS at 37°C for 30 min, followed by mechanical trituration.

Digestive enzymes were then removed after centrifugation of the cells at 100 g for 5 min. Isolated DRG neurons were resuspended in warm DMEM/F-12 (1% penicillin streptomycin and 10% FBS) and plated onto glass coverslips coated with poly-L-lysine (MilliporeSigma, Cat. Nu. P4707) and laminin (MilliporeSigma, Cat. Nu. L2020).

For overnight incubation with palmitoyl-ccPA (Cayman Chemical, Cat Nu. 10010293), the DRG neurons were allowed to fully adhere to the slips (24 hours) before the media was replaced with fresh media or fresh media containing 300 µM palmitoyl-ccPA. DMEM/F-12 (1% P/S and 10% FBS) was used as the solvent for the palmitoyl-ccPA. The DRG neurons were incubated with palmitoyl-ccPA for 24 hours before whole-cell electrophysiology experiments. For experiments using FIPI, DRG neurons were used between 24-48 hours after isolation. The neurons were incubated in media containing 500 nM FIPI or the equivalent volume of DMSO (0.005%) for 30 minutes, and cells were patched in extracellular solution containing 500 nM FIPI.

Mechanically-activated currents from DRG neurons were categorized by their inactivation kinetics as rapid, intermediate, or slow based on the criteria from our previous publications^8,58,59^. We considered currents to be rapidly adapting if they fully inactivated before the end of the 200-ms mechanical stimulus. The inactivation time constants (tau) of these currents were 14.16 ± 5.12 ms (mean ± SD) similar to our previous work^8,58,59^. This inactivation time constant was very similar to PIEZO2 transiently transfected in N2A-P1KO cells (**Supplementary Fig. 3 a, b, e and Supplementary Fig. 4a**). Intermediately adapting currents did not fully inactivate but the leftover current at the end of the mechanical stimulation was <50% of the peak current. The inactivation time constant for these currents was 30.67 ± 10.70 ms. Slowly adapting currents also did not fully inactivate, and the leftover current at the end of the mechanical stimulation was >50% of the peak current. The inactivation constant for these currents was 111.85 ± 52.47 ms.

### hiPSC-Derived Sensory Neurons

hiPSC-derived sensory neurons (RealDRG^TM^, Anatomic Incorporated, Cat. Nu. 1020F1-1M) were cultured and differentiated according to manufacturers (Anatomic Incorporated) instructions. Briefly, hiPSC cells stored in liquid N_2_ were thawed and plated onto poly-L-ornithine (MilliporeSigma, Cat. Nu. P4967) and iMatrix-511 SILK (Anatomic Incorporated, Cat. Nu. M511S) coated 12 mm coverslips. Cells were cultured in differentiation Chrono Senso-MM media (Anatomic Incorporated, Cat Nu. 7008) with no antibiotics present for 12-16 days. Media was changed every 48 hours. The hiPSC derived neurons were incubated in media containing 500 nM FIPI or the equivalent volume of DMSO (0.005%) for 30 minutes and were patched in extracellular solution containing 500 nM FIPI.

### TIRF Microscopy

N2A-P1KO cells were transiently transfected with cDNA encoding *Piezo2-GFP* and *Tmem120a-tdTomato*, *Tmem120a-* W193A*-tdTomato*, *Tmem120a-tdTomato* (AxxAA) or *tdTomato* (500 ng / construct, Effectene). The next day, the cells were plated onto poly-L-lysine coated 350mm round coverslips (#1.5 thickness; ThermoFisher Scientific). The cells were used for TIRF imaging two days after transfection. Coverslips were placed into a recording chamber filled with extracellular solution (137 mM NaCl, 5 mM KCl, 1 mM MgCl_2_, 2 mM CaCl_2_, 10 mM HEPES, and 10 mM glucose, pH adjusted to 7.4 with NaOH). TIRF images were obtained at room temperature using a Nikon Eclipse Ti2 microscope. Fluorescence excitation was performed using a 15 mW solid state 488 and 561 nm laser at 90% of the maximal power through a CFI Aprochromat TIRF 60X oil objective (N.A.: 1.49). Images were captured using an ORCA-Fusion Digital CMOS camera (Hamamatsu) through emission filters 525/50 and 600/50 nm for the green and red channels, respectively.

The images were analyzed using Nikon NIS-Elements AR Analysis software. Regions of interest (ROIs) were generated by hand drawing the outline of individual cells allowing for analysis of fluorescent signal over the total rea of each cell in the TIRF angle. Mean fluorescence intensities within these single-cell ROIs were averaged for each coverslip (5-32 total cells / coverslip) and plotted.

### Behavioral Testing

Experiments were performed on male and female mice (C57Bl/6J, The Jackson laboratory, Bar Harbor, Maine) (age 5–8 weeks; weight 22–25 g). Mice were kept in the Laboratory Animal Housing Faculty of Rutgers New Jersey Medical School at 24–25°C, provided with standard mouse chow and water *ad libitum* and maintained under a 12 h light/dark cycle. Animals were acclimatized to the testing room for at least 1 h before all behavioral tests. The same experimenter handled and tested all animals in each experiment and was blinded to the treatment. We used both male and female mice, and the data pooled together as we did not observe a significant difference between the sexes. FIPI (in DMSO stock) was diluted in mixture of Kolliphor EL (Sigma-Aldrich, Cat Nu. C5135) and standard PBS (Gibco) in the ratio 1:1:48 with final concentration of FIPI 0.5 mg/ml. Mice were intraperitoneally injected with FIPI (n=14) or vehicle (n=14) at the dose 3 mg/kg with a 0.5 ml Hamilton syringe and 30 g needle and returned in the home cage for 2 hours. After drug administration, mice were allowed to acclimatize on a metal mesh platform in an individual Plexiglas chamber for 1 hour.

Mechanical sensitivity was assessed by measuring paw withdrawal thresholds to von Frey filament (Stoelting, Wood Dale, IL) stimuli. For determining the 50% withdrawal threshold using the Up-Down method, calibrated von Frey filaments were applied as previously described^60,61^ and found in the von Frey Test protocol (Standard Operating Procedure The Jackson Laboratory Mouse Facility). Briefly, testing initiated by application filaments to the soft pad of the hind paw between the tori at the base of the digits for 2-3 sec using the filament force in the mid-range (0.4 g). After positive response (defined as withdrawal, shaking, splaying, lifting or licking of the paw), we apply the next lower von Frey filament. If the animal did not respond, we apply the next higher von Frey filament. Four additional measures are recorded after the first change in response for both left and right paw. The 50% threshold is then calculated using the formula^61^: 50% threshold (g) = 10(X+kd), where X = log10 (target force in grams of the final von Frey filament used), k = tabular value for the response pattern (see Appendix 1 in reference^60^ and d = the average increment (in log units) between the target force in grams of the von Frey filaments used (2.0, 1.4, 1.0, 0.6, 0.4, 0.16, 0.07, 0.04 g).

Thermal sensitivity of the paw was assessed with a Model 336 Analgesia Meter (IITC Life Science) as described previously^62^. After performing von Frey mechanical testing mice were transferred and acclimatized on a glass floor in same Plexiglas chambers for 30 min, after which a light beam was focused on the midplantar region of the hind paw. The latency to respond by withdrawing the paw from the light was recorded. The cutoff time was set up to 20 s to prevent tissue damage. Stimuli were applied to the paws three times at 5 min intervals, and the average latency was calculated^62^.

The experimenter for all behavioral experiments was blind to what treatment each mouse received during testing and quantification of the data.

### cDNA Constructs

The GFP-tagged *Piezo1* construct (PIEZO1-GFP) was generated by subcloning the mouse *Piezo1* to the pcDNA3.1(-) vector from the original IRES-GFP vector then PCR cloning GFP and ligating it to the N-terminus of *Piezo1*^37^. The GFP-tagged *Piezo2* construct (PIEZO2-GFP) was generated by PCR cloning GFP and ligating it to the N-terminus of *Piezo2* in the pCMV SPORT6 vector. The tdTomato-tagged *Tmem120a* construct was generated by PCR cloning *Tmem120a* using the Origene MR205146 clone as a template and subcloning it to the ptdTomato-N1 vector (Clontech), placing the tdTomato tag to the C-terminus of *Tmem120a*^8^. For PCR cloning, the Pfu-Ultra proofreading enzyme (Agilent) was used and the constructs were verified with sequencing. The *Tmem120b* cDNA clone was purchased from Origene (MR205067, NM_001039723). The Opto-PLD active and inactive constructs were purchased from Addgene (140114, and 140061). The hPLD1 cDNA clone was a kind gift from Dr.

Michael A. Frohman (Genbank accession number = U3845). The mPLD2 cDNA clone was a kind gift from Dr. Guangwei Du (Genbank accession number = 87557). Point mutations of *Tmem120a* were generated using the QuikChange II XL Site Directed Mutagenesis Kit (Agilent, Cat. Nu. 200522) per the manufacturer’s instructions. Full plasmid sequencing (Plasmidsaurus) was used to confirm that the intended residues were mutated and that no off-target mutations were present.

### Statistics

Data are represented as mean ± SEM plus scatter plots as indicated in figure legends. To determine whether data significantly deviated from normal distribution, Shapiro-Wilk tests were used. To determine whether data between groups had a significant difference in variance, Levene’s test for homogeneity of variance (centered on median) was used. When comparing between groups for data that did not significantly deviate from a normal distribution or variance, statistical significance was calculated either with a two-sample *t* test (two tailed), ANOVA with post-hoc Tukey. And for between group comparisons with data that significantly deviated from normality, Mann-Whitney or Kruskal-Wallis tests were used. When comparing within groups with repeated measures, a repeated-measures ANOVA & paired-sample t-test for normally distributed data. To compare nominal data (distribution of inactivation kinetics from mouse DRG neurons), a Chi-Square test was used. When multiple comparisons were necessary, the threshold for significance was corrected for the number of comparisons (α = 0.05 / # of comparisons). The specific tests and sample sizes for each for each experiment are described in the figure legends and noted in the figures. All statistical testing was performed using R Commander. Data plotting was performed using the Origin 2021 software.

## Data Availability

Data will be made available upon request to the corresponding author.

## ACKNOWLEDGEMENTS

This study was supported by National Institutes of Health grant R01-NS055159 to T. R. and by NCI-CCSG P30CA072720-5923 to the Rutgers Cancer Institute of New Jersey Metabolomics Shared Resource Metabolomics facility. The human PLD1 clone was a kind gift from Dr. Michael A. Frohman (Stony Brook University). The mouse PLD2 clone was generously provided by Dr. Guangwei Du (UTHealth, Houston). The authors are grateful to Patrick Walsh and Vincent Truong from Anatomic Incorporated for providing the hiPSC-derived human peripheral sensory neurons.

## AUTHOR CONTRIBUTIONS

M.G. and T.R conceptualize the study. M.G. performed all electrophysiology and TIRF experiments and the LC-MS/MS sample preparation. Y.Y. performed the behavioral experiments. Y.W. performed and X.S. supervised the the LC-MS/MS experiments.

M.G. visualized the data. T.R. supervised the study. T.R. and M.G. wrote the manuscript, all authors reviewed and revised the manuscript.

---

**Supplementary Fig. 1.**
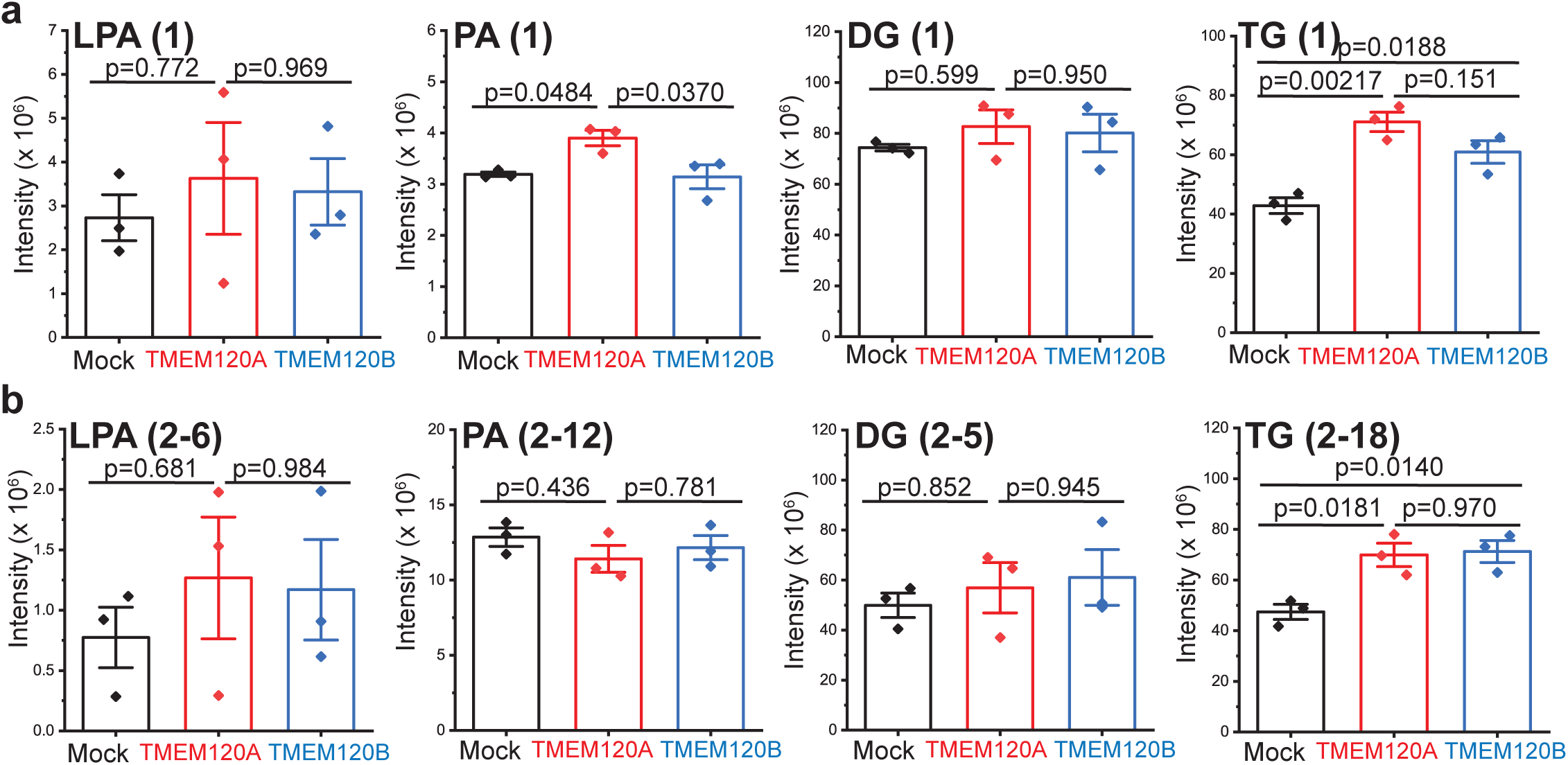
TMEM120A expression does not substantially change levels of unsaturated lipids. LC-MS/MS experiments using N2A-Pz1-KO cells transiently transfected with vector (mock, black), TMEM120A (red), or TMEM120B (blue) as described in the methods section. **(a)** Scatter plots and mean ± SEM of the relative mono-unsaturated lipid intensities detected for n=3 transfections of LPA (ANOVA, F(2,6)=0.253, p=0.784), PA (ANOVA, F(2,6)=6.872, p=0.0281), DG (ANOVA, F(2,6)=0.533, p=0.612), TG (ANOVA, F(2,6)=18.98, p=0.00254). Post-hoc Tukey tests displayed on plots. **(b)** Scatter plots and mean ± SEM of the relative poly-unsaturated lipid intensities detected for n=3 transfections of LPA (ANOVA, F(2,6)=0.417, p=0.677), PA (ANOVA, F(2,6)=0.869, p=0.466), DG (ANOVA, F(2,6)=0.382, p=0.698), TG (ANOVA, F(2,6)=10.92, p=0.01). Post-hoc Tukey tests displayed on plots.

**Supplementary Fig. 2.**
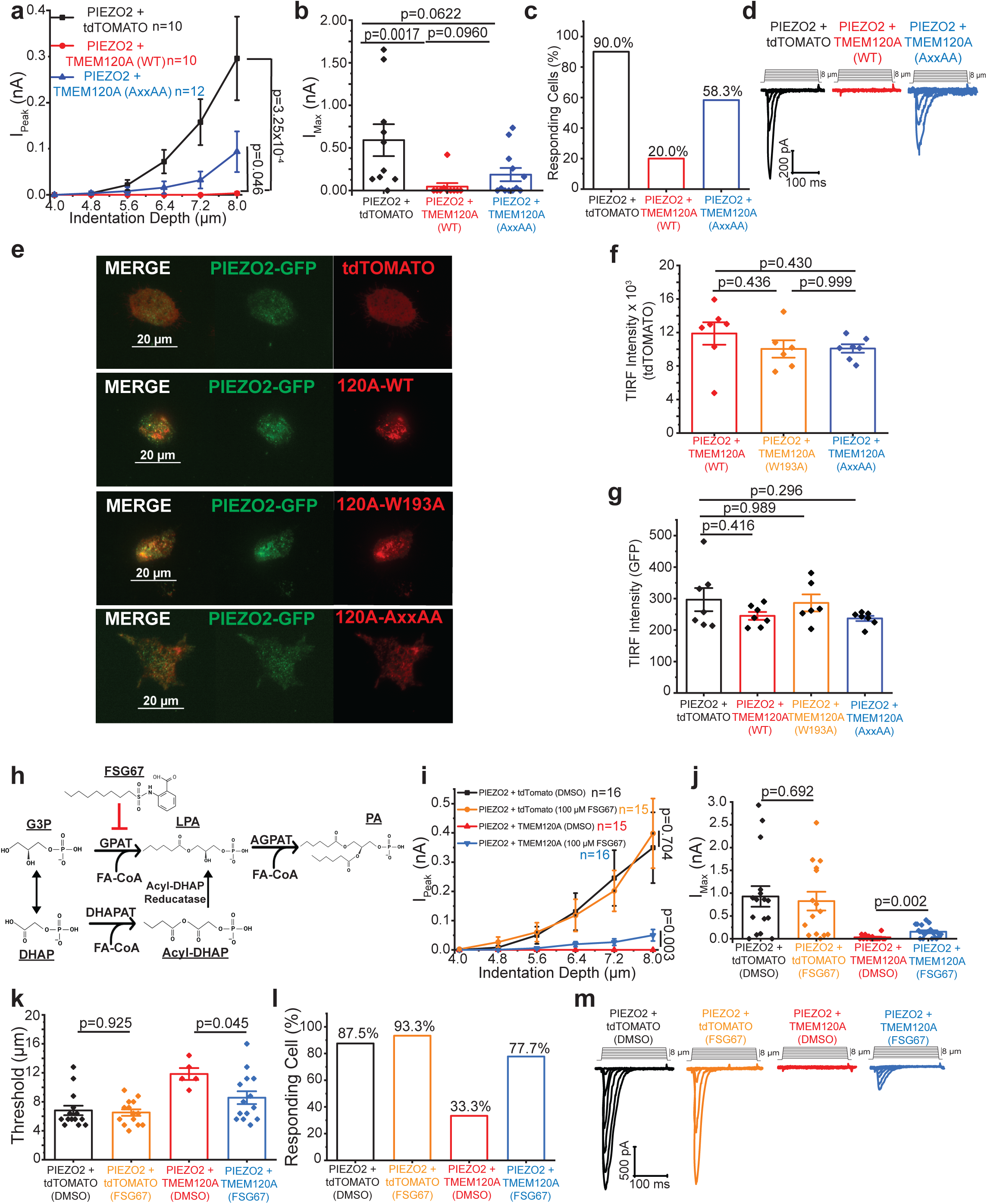
TMEM120A dependent changes in cellular lipid content contribute to PIEZO2 inhibition. **(a)** Whole-cell patch clamp experiments at -60 mV in cells transiently transfected with PIEZO2-GFP with tdTOMATO (black), TMEM120A-WT-tdTom (red), or TMEM120A-AxxAA-tdTom (AxxAA = W193A, H196A, H197A; blue). Currents amplitudes are plotted (mean ± SEM) and statistical difference of AUC for 4.0-8.0 µm stimuli calculated with the Mann-Whitney test. **(b)** Scatter and mean ± SEM for maximum currents (Kruskal-Wallis, χ2=10.769, df=2, p=0.004; Mann-Whitney tests displayed). **(c)** Percent of cells displaying mechanically-activated currents. **(d)** Representative current traces. **(e)** TIRF microscopy was performed as described in the methods section. Representative images (“120A” = TMEM120A). **(f)** Scatter and mean ± SEM for the fluorescence intensity for TMEM120A-WT-tdTom, TMEM120A-W193A-tdTom, and TMEM120A-AxxAA-tdTom co-expressed with PIEZO2-GFP (ANOVA, F(2,17)=1.08, p=0.362; post-hoc Tukey tests displayed). **(g)** Scatter and mean ± SEM for the fluorescent intensity of PIEZO2-GFP co-expressed with tdTOMATO, TMEM120A-WT-tdTom, TMEM120A-W193A-tdTom, or TMEM120A-AxxAA-tdTom (ANOVA, F(3,23)=1.571, p=0.224; post-hoc Tukey tests displayed). **(h)** Scheme of LPA and PA synthesis and GPAT inhibition by FSG67. **(i)** Whole-cell patch clamp experiments at -60 mV in cells transiently transfected with PIEZO2-GFP with tdTOMATO (black, DMSO-treated; orange, FSG67-treated) or TMEM120A-tdTom (red, DMSO-treated; blue, FSG67-treated). Current amplitudes are plotted (mean ± SEM) and statistical difference of AUC for 4.0-8.0 µm stimuli calculated with the Mann-Whitney test. **(j)** Scatter and mean ± SEM for maximum currents (Kruskal-Wallis, χ2=26.78, df=3, p=6.53x10-6; Mann-Whitney tests displayed). **(k)** Scatter and mean ± SEM for the threshold of membrane indentation to elicit mechanically-activated-current (Kruskal-Wallis, χ2=12.43, df=3, p=0.006; Mann-Whitney tests displayed). **(l)** Percent of cells displaying mechanically-activated currents. **(m)** Representative current traces.

**Supplementary Fig. 3.**
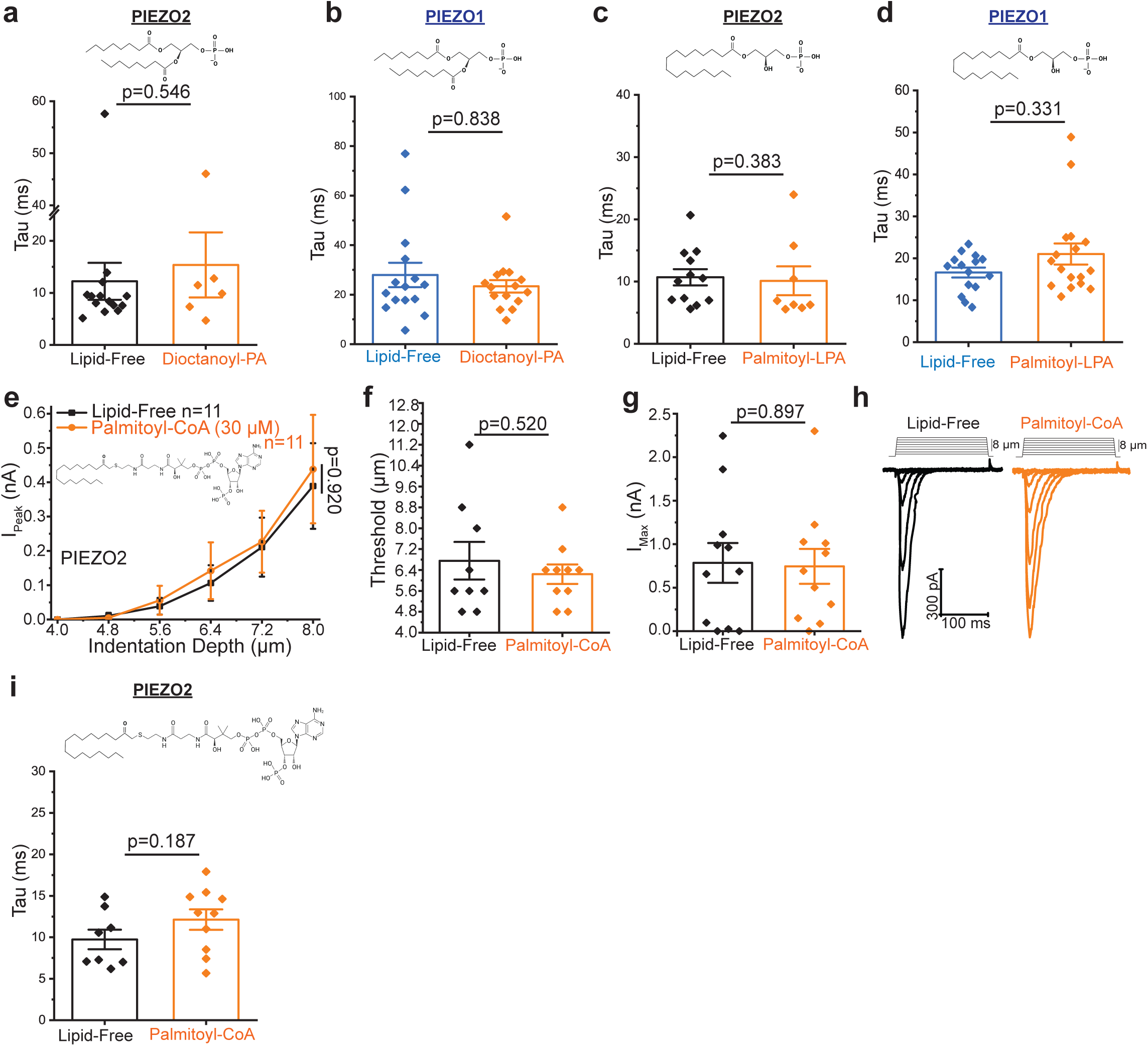
PA, LPA and CoA do not change the inactivation kinetics of PIEZO2 or PIEZO1, and CoA does not inhibit PIEZO2. The inactivation time constants (tau) are calculated as described in the methods section. **(a)** PIEZO2-GFP transfected cells supplemented with dioctanoyl-PA (Mann- .Whitney) **(b)** PIEZO1-GFP transfected cells supplemented with dioctanoyl-PA (Mann-Whitney). **(c)** PIEZO2-GFP transfected cells supplemented with palmitoyl-LPA (Mann-Whitney). **(d)** PIEZO1-GFP transfected cells supplemented with palmitoyl-LPA (Mann-Whitney). **(e-i)** Whole-cell patch-clamp experiments at -60 mV in N2A-Pz1-KO cells transiently transfected with PIEZO2-GFP with patch-pipette solution supplemented with palmitoyl-CoA as described in the methods section. **(e)** Current amplitudes are plotted (mean ± SEM) and statistical difference of AUC 4.0-8.0 µm stimuli calculated with the Mann-Whitney test. **(f)** Membrane indentation depth threshold to elicit mechanically-activated currents (t-test). **(g)** Maximum current amplitudes (t-test). **(h)** Representative current traces. **(i)** Inactivation constants (tau) for lipid-free and palmitoyl-CoA treated cells (t-test).

**Supplementary Fig. 4.**
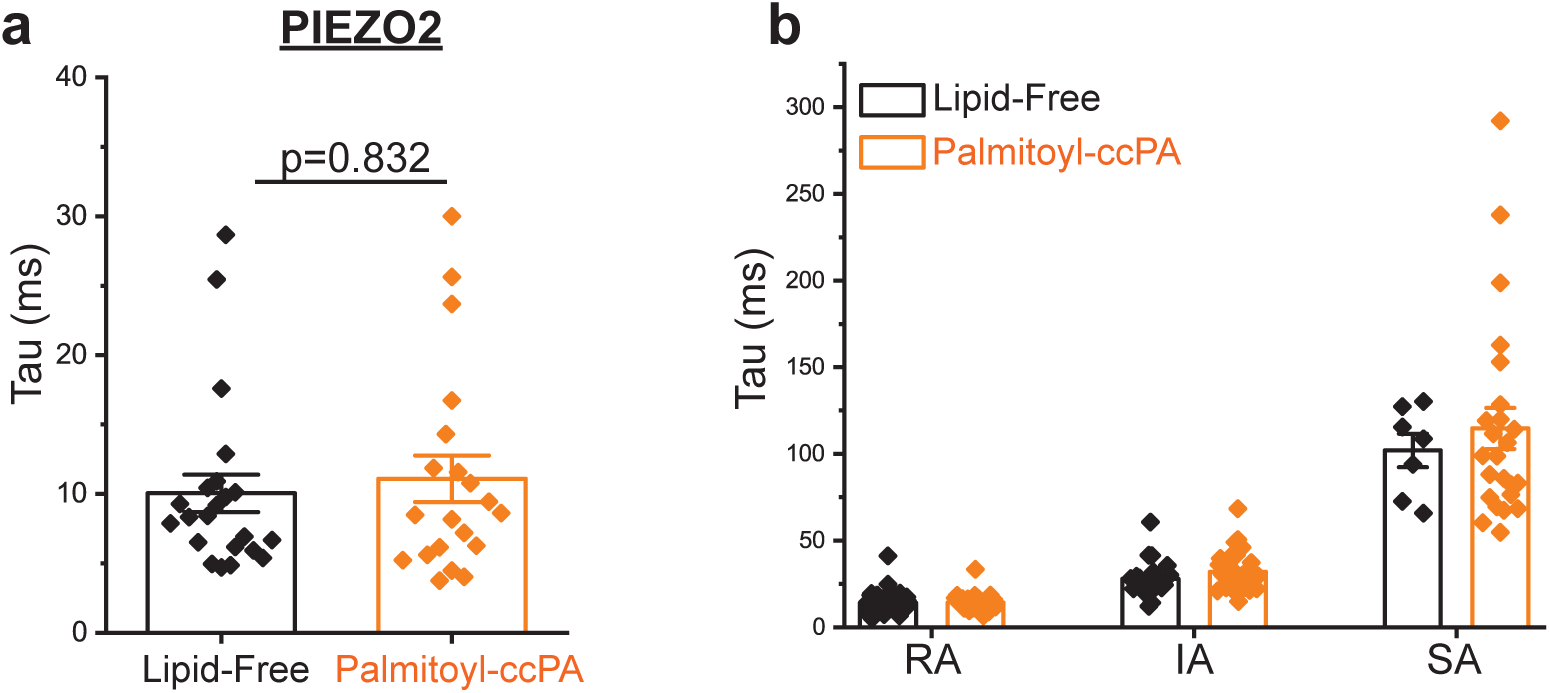
Extracellular application of ccPA does not change the inactivation kinetics of PIEZO2. The inactivation time constants (tau) are calculated as described in the methods section. **(a)** PIEZO2-GFP transfected cells treated with palmitoyl-ccPA (Mann-Whitney). **(b)** Isolated mouse DRG neurons treated with palmitoyl-ccPA.

**Supplementary Fig. 5.**
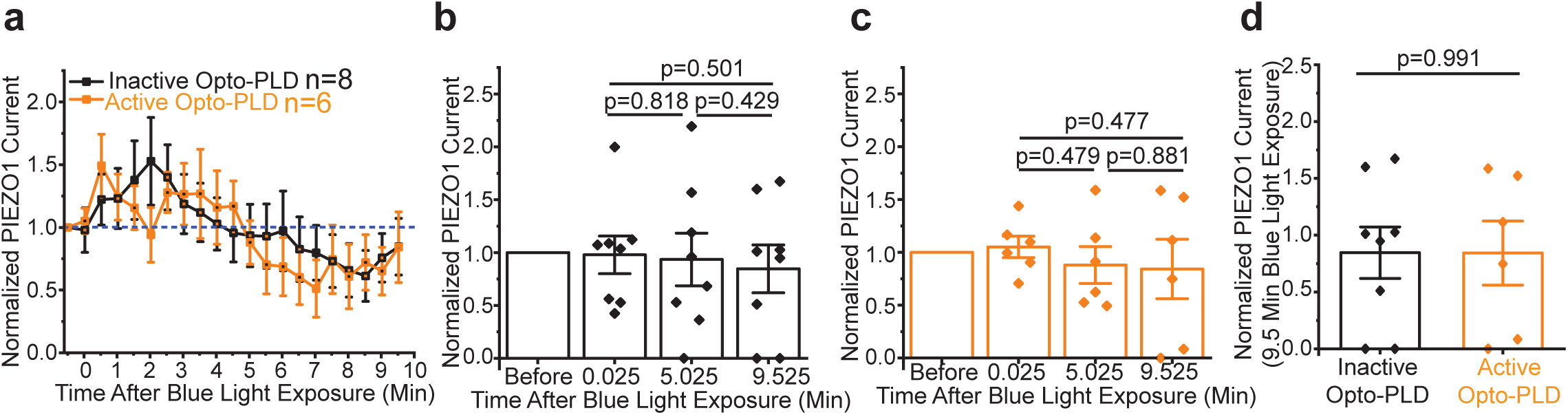
Optogenetic activation of PLD does not inhibit PIEZO1. Whole-cell patch-clamp experiments at -60 mV in N2A-Pz1-KO cells transfected with PIEZO1-GFP and active Opto-PLD (black) or inactive Opto-PLD (orange; H170A) as described in the methods section. **(a)** PIEZO1-GFP currents with fixed, continuous membrane indentations after blue light exposure normalized to currents before blue light. **(b)** Scatter and mean ± SEM for PIEZO1-GFP co-expressed with inactive Opto-PLD after blue light exposure (Repeated-Measures ANOVA, F=0.344, df=2,14, p=0.714; paired t-tests displayed). **(c)** Scatter and mean ± SEM for PIEZO1-GFP co-expressed with active Opto-PLD after blue light exposure (Repeated-Measures ANOVA, F=0.364, df=2,10, p=0.703; paired t-tests displayed). **(d)** Scatter and mean ± SEM of PIEZO2-GFP with inactive or active Opto-PLD after 9.5 min of blue light exposure (t-test).

**Supplementary Fig. 6.**
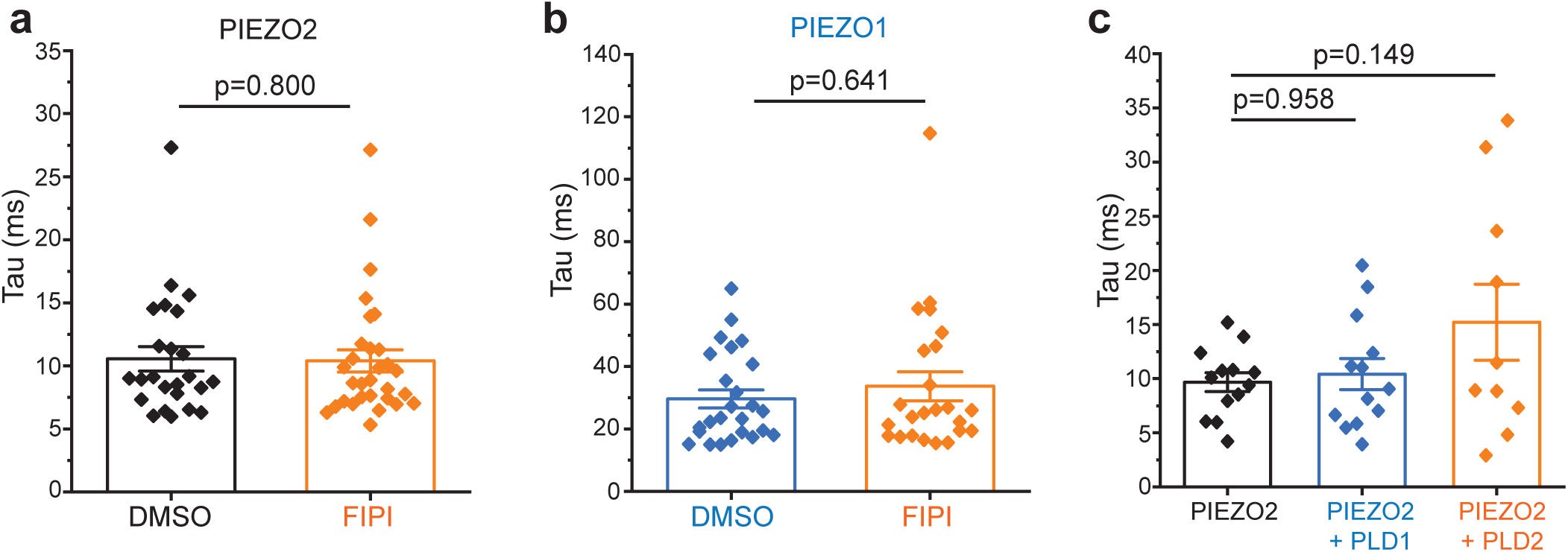
Phospholipase D (PLD) enzymes do not affect the inactivation kinetics of PIEZO1 or PIEZO2. The inactivation time constants (tau) are calculated as described in the methods section. **(a)** PIEZO2-GFP transfected cells treated with FIPI or equivalent volume of DMSO (Mann-Whitney). **(b)** PIEZO1-GFP transfected cells treated with FIPI or equivalent volume of DMSO (Mann-Whitney). **(c)** N2A-Pz1-KO cells transiently transfected with PIEZO2-GFP alone (black), PLD1 (blue), PLD2 (orange)(ANOVA, F=2.085, df=2,33, p=0.140; post-hoc Tukey tests displayed).

**Supplementary Fig. 7.**
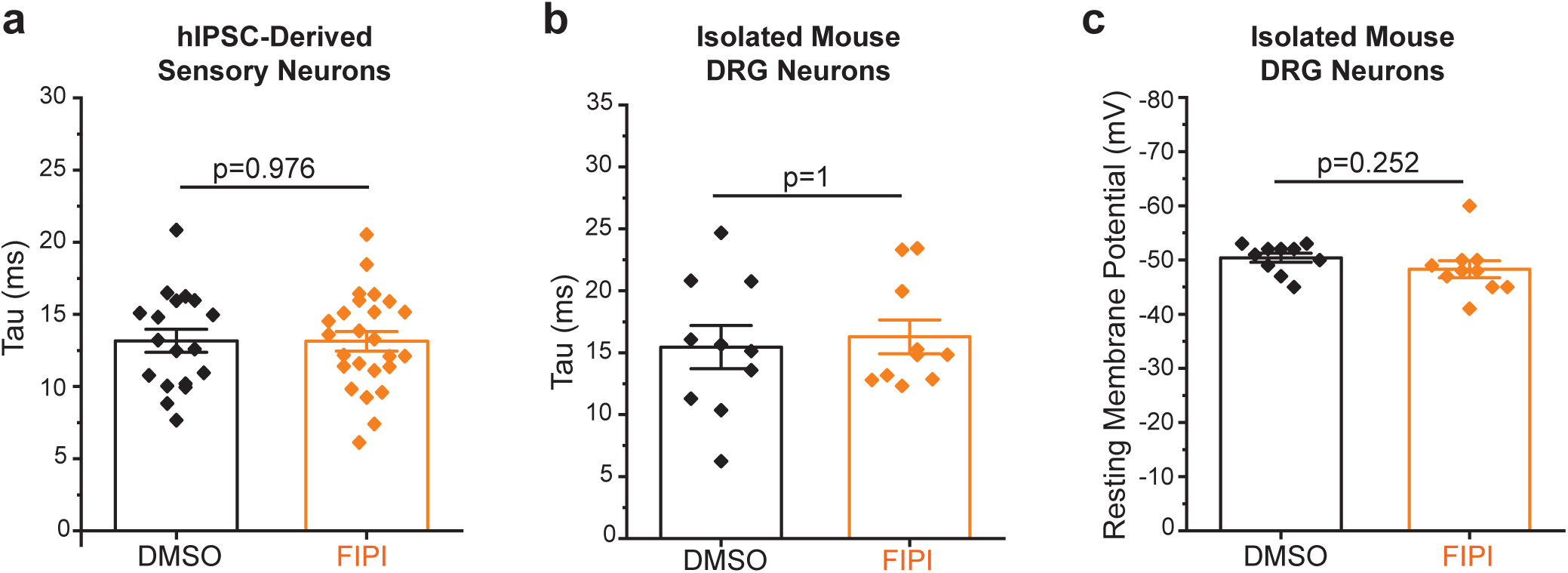
Phospholipase D (PLD) inhibition does not affect endogenously expressed PIEZO2 inactivation kinetics or the resting membrane potential of DRG neurons. The inactivation time constants (tau) are calculated as described in the methods section. **(a)** hiPSC-derived sensory neurons treated with FIPI or equivalent volume of DMSO (Mann-Whitney). **(b)** Isolated mouse DRG neurons treated with FIPI or equivalent volume of DMSO displaying RA type mechanically-activated current (Mann-Whitney). **(c)** Resting membrane potential of isolated mouse DRG neurons treated with FIPI or equivalent volume of DMSO (t-test).

